# Exercise-induced differential transcriptional output of AMPK signalling improves axon regeneration and functional recovery

**DOI:** 10.1101/2024.12.08.627390

**Authors:** Sibaram Behera, Anindya Ghosh Roy

## Abstract

In adulthood, the regenerative capacity of the injured brain circuit is poor preventing functional restoration. Rehabilitative physical exercise is a promising approach to overcome such functional impairment and the metabolic sensor AMPK has emerged to be a critical mediator for this. However, the mechanistic understanding of upstream and downstream components of AMPK signalling in the physical exercise-mediated enhancement of axon regeneration is not clear. We combined swimming exercise with laser axotomy of posterior lateral microtubule (PLM) neurons of *Caenorhabditis elegans* to address this question. We found that direct activation of AMPK through AICAR treatment is sufficient to improve axon regeneration and functional recovery. The PAR-4/Liver kinase B1 (LKB1) acts upstream of AMPK to enhance functional recovery following swimming exercise. Using genetics, tissue-specific RNAi, and AICAR treatment, we found that the transcriptional regulators DAF-16 and MDT-15 act downstream of AMPK in mediating the positive effects of swimming. We found that MDT-15 acts in neuron to mediate the benefit of AMPK activation in axon regeneration, whereas DAF-16 acts both in neuron and muscle to promote regrowth downstream to AMPK. We also showed that swimming exercise induces nuclear localization of DAF-16 in an AMPK-dependent manner. Our results showed that neuronal and non-neuronal arms of AMPK signalling play an integrative role in response to physical exercise to promote functional recovery after axon injury.

**Significance statement:** Finding ways to promote functional recovery after accidental damage to the nervous system has been challenging as adult neurons lose the capability to regenerate. Rehabilitation therapy is the most promising approach to improve the health condition of patients with nervous system injury. Even in the roundworm *C. elegans*, axon regeneration could be enhanced through swimming exercise, which is mediated by the metabolic energy sensor AMP Kinase. In this study, using sensory neurons in worm, we found that PAR-4/ Liver kinase B1 acts upstream of AMPK. Whereas, the transcription factor DAF-16/ FOXO and the transcriptional co-regulator MDT-15 act as downstream signalling arms in muscle and neuron tissues. Excitingly, both of these arms could be harnessed through agonist-mediated activation of AMPK to promote functional recovery in adulthood.

## Introduction

Central nervous system (CNS) injury like traumatic brain injury (TBI) and spinal cord injury (SCI) is a growing socio-economical challenge due to their devastating consequences on quality of life (1, 2). Injury to the CNS axons hamper information transfer resulting in functional impairment because of axon regeneration failure due to extrinsic and neuron-intrinsic factors (3, 4). In the model systems of axon regeneration study, the manipulation of a single regulator can only provide limited functional recovery, which demands a combinatorial strategy for achieving desired functional recovery (3). Clinically validated rehabilitative therapy involving physical exercise is promising for functional recovery and can be combined with other regeneration-enhancing strategies (5). Physical exercise has well-documented beneficial effects on health parameters such as improvement in motor function, cognitive function, memory and neurogenesis in humans (6–8). Physical exercise also leads to improved functional recovery and axon regeneration following injury in various model systems (9–11). However, the cellular and molecular mechanism of how physical exercise improves axon regeneration and functional recovery is still unclear.

The metabolic sensor AMPK (AMP-activated protein kinase) is a key mediator of the positive effects of exercise(12) including axon regeneration (10, 12). It acts as a cellular energy sensor and regulates energy homeostasis by monitoring the ATP level through the competitive AMP/ATP binding in the γ subunit that regulates its activity (13). AMPK is also essential for swimming exercise-mediated improvement in behavioural recovery, fitness preservation and muscle health (10, 14, 15). Phosphorylation of Thr172 in the α subunit is critical for AMPK activity (16) and it is mediated by the upstream kinase Liver Kinase B1 (LKB1) and CAMKK2 (also known as CAMKKβ) (17–21). The activated AMPK normally inhibits the anabolic processes while activating catabolic processes to maintain energy homeostasis by direct phosphorylation of key metabolic enzymes and also by regulating several transcription factors for restoring energy balance (22, 23). How AMPK signalling regulates axon regeneration, especially following a physical exercise paradigm is not well understood. Emerging evidence of the tissue-specific role of AMPK and the cross-tissue interaction of this signalling demands further mechanistic understanding (24, 25).

*Caenorhabditis elegans* is an excellent model system to address this gap as it has been a well-established axon regeneration model for the last two decades (26, 27). Also, a physical exercise paradigm involving swimming mimics major features of mammalian exercise in worm (28, 29). Regeneration studies in multiple neurons including gentle touch sensory neuron and motor neuron has revealed positive and negative regulators of axon regeneration (27, 30). Calcium and DLK-1 MAP kinase signalling promotes axon regeneration while guidance factors and *let-7* microRNA inhibit axon regeneration (31–37). Axon regeneration in the early stages of life in both motor and touch neuron leads to functional recovery. However, functional rewiring is strongly inhibited in older age (37–40). Interestingly, swimming exercise following axotomy in PLM neuron promote axon regrowth and functional restoration in older age (10). The beneficial effect of swimming in axon regeneration depends on the metabolic sensor AMPK/AAK-2 (10).

In this study, we utilised the combined paradigm of laser-induced PLM neuron axon injury, swimming exercise, genetics and pharmacology to identify the upstream and downstream components of AMPK signalling in the improvement of axon regeneration through swimming exercise. We established that acute activation of AMPK using AICAR treatment mimics the effect of exercise for improving behavioural recovery. Further, we found that LKB1/PAR-4 is a key upstream molecule activating AMPK after swimming exercise. Our results also show that FOXO/DAF-16 and MDT-15/PGC1α- are key downstream transcriptional mediators through which AMPK promotes functional rewiring after axotomy. Nuclear localization of DAF-16 is enhanced following a swimming session in an AAK-2-dependent manner. Further, using tissue-specific RNAi, we demonstrated that DAF-16 is required in both muscle and neuron to regulate rewiring events while MDT-15 is required only in neurons. Our study revealed neuron-specific and non-neuronal integrative roles of AMPK signalling arms in physical exercise-mediated enhancement of axon regeneration.

## Results

### AMPK activation through AICAR treatment improves functional recovery after laser-induced PLM axon injury

Following an axonal injury, physical exercise improves functional recovery and axon regeneration (9, 10). A single session of 90-minute swimming improves functional recovery following laser axotomy of *C. elegans* posterior gentle touch sensory (PLM) neurons (Fig. 1A-B) (10). The metabolic sensor AMPK/AAK-2 (AMPK α subunit) is essential for this improvement of functional restoration due to swimming exercise (Fig.1B-C) as seen before (Kumar et al 2021). However, whether the direct activation of AMPK would replace the swimming session itself to promote functional recovery is not clear. The functional recovery is assessed in terms of the ‘recovery index’ which is determined by the ratio of the Posterior Touch Response Index (PTRI) value obtained at 24 h to the PTRI value at 3h post-axotomy (10, 37). We used AICAR as an agonist reported to directly activate AMPK (41, 42) and also previously used in *C. elegans* (43). The temporary paralysis of worms during a swimming session by brief levamisole exposure before the session begins blocks the beneficial effects of exercise (Fig. 1D) (10). We tested three concentrations of acute AICAR treatment in this set-up and observed that 1 mM AICAR treatment improved the recovery index significantly at the L4 stage (Fig. 1D-E). However, such enhancement in functional recovery is not observed in *aak-2(lf)* animals suggesting that the enhancement occurs through AMPK activation (Fig.1E). The recovery index is significantly reduced in day 3 adult (A3) axon injury (Fig. 1E) as seen before (37). Interestingly, acute AICAR treatment could overcome this age-related decline in recovery index at A3 stage axotomy experiments (Fig.1E). AICAR treatment also improved regrowth length at 24 h post-axotomy signifying enhanced axon regeneration process (Fig. 1F). Similarly, in the background with a constitutively active AAK-2 (CA-aak-2) mutant (44), the functional recovery after axon injury is significantly enhanced as compared to wildtype both in L4 and adulthood day 3 stage (Fig.1G).

**Figure 1.**
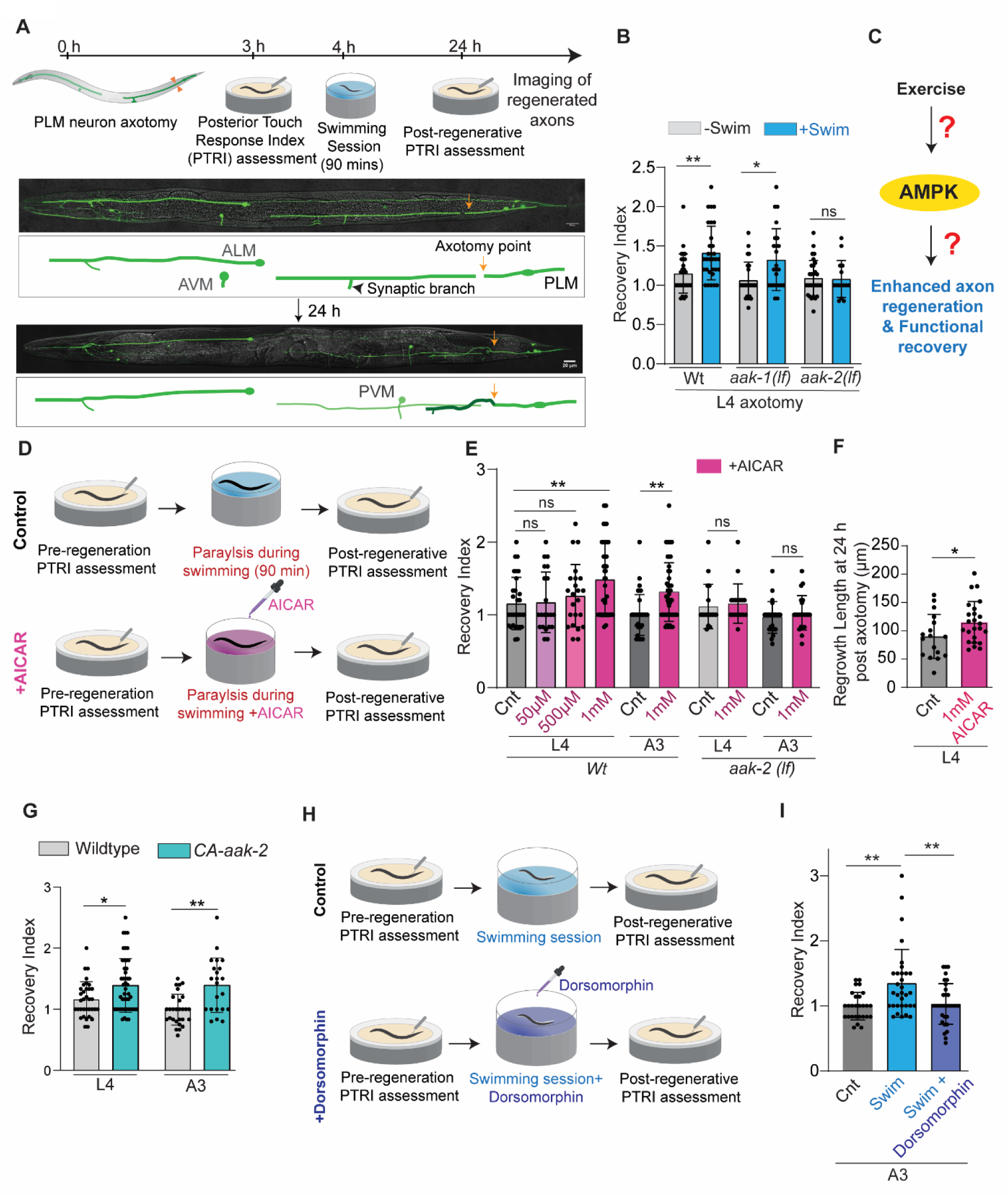
Direct activation of AMPK through acute treatment of AICAR improves axon regeneration**. (A)** Schematics of axon injury paradigm of the posterior gentle touch sensory neuron (PLM) in combination with swimming exercise. After a laser-induced break is introduced to the PLM axon, a posterior touch response assay was performed to record the posterior touch response index (PTRI) followed by a 90-minute swimming session. At 24 h of axotomy, PTRI is assessed followed by imaging of the regeneration. **(B)** Functional recovery is represented by the recovery index for *wildtype,aak-1(lf) and aak-2(lf)* animals. Axotomy was performed at the L4 stage. The recovery index is the ratio of PTRI at 24h post-axotomy to PTRI at 3h postaxotomy. n=15-31 number of worms, N=3-4 independent replicates. **(C)** Schematics representing uncharacterized AMPK signalling in the context of exercise and axon regeneration. **(D)** Schematics showing the AICAR treatment paradigm to transiently paralyzed worms during the swimming session. **(E)** Bar graph showing the effect of AICAR treatment on recovery index. N=3-4, n=18-55. **(F)** Axon regrowth characterization at 24h postaxotomy shows an enhancement following 1mM AICAR treatment. N=3,n= 17-24 **(G)** Recovery index shows an enhancement in behavioural recovery for constitutively active AMPK (CA-aak2) worms. N=3-4, n=22-47.**(H)** Schematics showing the dorsomorphin (Compound C) treatment paradigm. **(I)** Dorsomorphin treatment (10 µM) significantly reduced the enhanced recovery mediated by swimming. N=3-4, n=27-33. Statistics, for B, E, G, I *p < 0.05, **p < 0.01 ***p < 0.001 ANOVA with Tukey’s multiple comparison test. For F *p < 0.05 unpaired t-test. Error bars represent SD; ns, not significant.

Conversely, treatment with Dorsomorphin (Compound C), which is a well-known inhibitor of AMPK activity (45, 46) during the swimming session (Fig. 1H) blocked the enhanced functional recovery due to the swimming session, validating AMPK activation as a key mediator of swimming exercise benefits (Fig. 1I).

We wanted to test the effect of AICAR treatment in an alternate functional recovery paradigm, which also utilizes the AMPK function. For example, the Posterior touch response index decreases significantly as the worm ages from day-4 adult stage (A4) to day-5 adult stage (A5) and when the worms are subjected to a 90-minute swimming session at A4 stage this decline was not observed (Fig. S1A) as seen before (10, 37). AICAR treatment to temporarily paralyzed A4 stage worms resulted in the preservation of posterior gentle touch response in the A5 stage (Fig. S1B-C). Also, brief treatment of dorsomorphin during the swimming session hinders the preservation of PTRI conferred by swimming exercise (Fig.S1D-E). These results established AMPK as a key regulator of exercise’s effect on axon regeneration and a promising therapeutic target.

### Liver kinase B1/PAR-4 acts upstream of AMPK in swimming-mediated enhancement of axon regeneration

We wanted to identify the enzyme that acts upstream of AMKP during swimming exercise, that promotes axon regeneration (Fig.2A). The AMPK enzyme is phosphorylated at the Threonine-172 position during physical exercise either by the liver kinase B1 or by CAMKK2 (17–21). In worms, the orthologues for liver kinase B1 and CAMKK2 are known as PAR-4 and CKK-1, respectively. The scaffolding protein PRY-1 or AXL-1, the worm orthologue of Axin is a key mediator of LKB-1 phosphorylation of AMPK (47–49).

**Figure 2.**
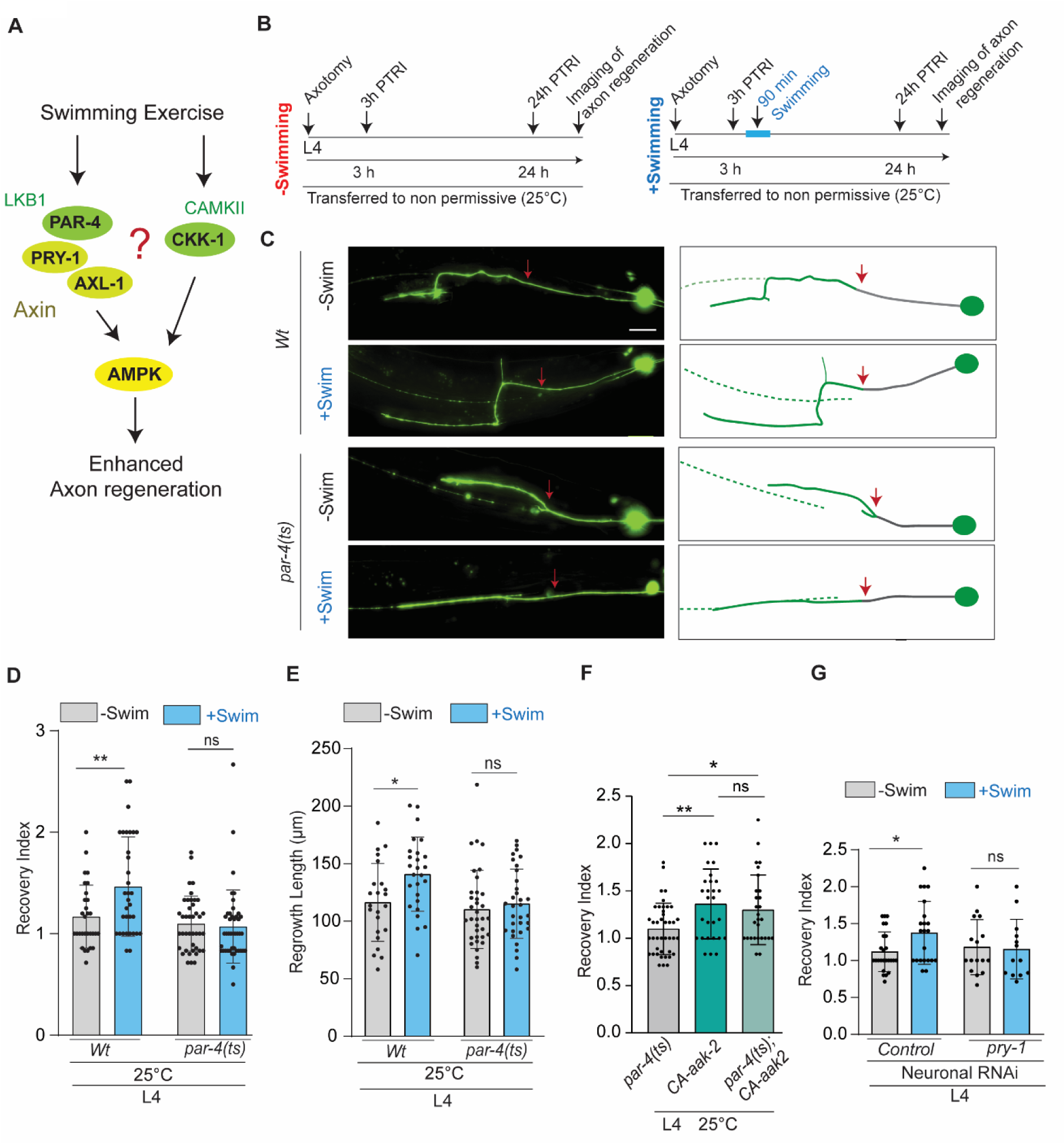
Swimming-mediated improvement in functional recovery after axon injury of PLM neuron requires liver kinase B1/PAR-4. **(A)** Schematics showing upstream regulators of AMPK that can be involved in swimming-mediated improvement in axon regeneration. **(B)** Paradigm to evaluate the role of PAR-4 using *par-4(ts)* mutants. The PLM axon was severed using a controlled laser pulse at the L4 stage and functional assessment was done by analysing the PTRI values. The worms were grown till the L4 stage at permissive temperature (15 °C) and after axotomy, were subjected to non-permissive temperature (25 °C). **(C)** Microscopic image showing the axonal regrowth at 24 h post axotomy with or without swimming exercise for wild type and *par-4(ts)* worms. The red arrow shows the position of the axotomy. Scale bar: 10 µm **(D)** Bar graph showing the recovery index values for wild type and *par-4(ts)* worms at 24 h post axotomy with the introduction of a 90-minute swimming exercise. N = 3-4 independent replicates, n = 30–43 number of worms. **(E)** Quantification of regrowth length at 24 h as represented in panel D.N=3-4,n=23-35. **(F)** Improved behavioural recovery mediated by constitutive activation of AMPK (CA-*aak2*) is not affected by *par-4(ts)*. N=3,n=28-41.**(G)** Effect of swimming exercise on the recovery index of neuronal knockdown of pry-1.N=2, n=13-25.Statistics, for D, E, F *p < 0.05, **p < 0.01 ANOVA with Tukey’s multiple comparison test. For G, *p < 0.05 Unpaired t-test Error bars represent SD; ns, not significant.

We found that in the absence of either *ckk-1* or *axl-1*, the recovery index is elevated significantly due to swimming exercise similar to the wild-type group (Fig.S2A-B). The regrowth at 24 h post-axotomy in these mutants is also comparable to wild-type condition (Fig.S2C), which ruled out the function of CKK-1 and AXL-1 in swimming exercise-mediated improvement in behavioural recovery. The aging paradigm of functional recovery also supported this finding as improved PTRI at day 5 adults was observed with swimming in *ckk-1* or *axl-1* mutant (Fig. S2D-E).

Since the absence of *par-4* leads to embryonic lethality (Kemphues et al., 1988), to test the role of PAR-4, we took advantage of a temperature-sensitive allele of *par-4*, which displays mutant phenotype when shifted to 25°C (Fig. 2B) (Watts et al., 2000). We saw a significant enhancement in the recovery index in the wild-type animals due to a swimming session at 25° C whereas such enhanced improvement was perturbed significantly in the *par-4(ts)* worms at 25°C temperature (Fig.2C). Although the basal level axon regeneration length is comparable to wild type (Fig.2D-E), swimming-mediated increment in regrowth length is not observed in *par-4(ts)* as opposed to wild-type condition (Fig.2D-E). In the assay for swimming-mediated improved touch sensation of aged worms, significant enhancement in PTRI of day 5 *par-4(ts)* animals was observed only at permissive (15°C) temperature but not at the non-permissive (25°C) temperature (Fig. S2F-G).To test whether *par-4* acts upstream of the activation of AMPK, we evaluated the effect of *par-4(ts)* in the background of constitutively active AAK-2. We found that constitutively active AAK-2 in the *par-4(ts)* background still shows an improved recovery at non-permissive 25°C at the comparable level to the active AAK-2 background (Fig.2F). However, activation of AMPK through AICAR treatment in the *par-4(ts)* mutant fails to mimic the exercise’s effect in axon regeneration (Fig.S2H-I). Also, only at 15° C, an improvement in Posterior Touch Response was observed in aged animals following AICAR treatment but not at 25° C (Fig.S2J-K).

This supports the role of PAR-4 in AICAR-mediated AMPK activation similar to previous reports (19). The knockdown of *pry-1* using RNAi perturbs swimming session-mediated enhancement in functional recovery following axotomy of PLM (Fig. 2G). These results showed that PAR-4/ LKB-1 and its scaffold PRY-1 are required for swimming exercise-mediated enhancement in sensory function in older age and also in behavioural recovery through axon regeneration.

### Transcriptional regulators, DAF-16 and MDT-15 mediate AMPK’s beneficial effects after swimming exercise

The activation of AMPK following physical exercise leads to an adaptive response by regulating transcription factors along with acute metabolic changes (23, 50, 51). AMPK directly phosphorylates the FOXO3/ DAF-16 transcription factor in mammals and *C.elegans* (52, 53). In mice, AMPK also directly phosphorylate the CREB family transcription factor and histone deacetylase HDAC5 (54–56). AMPK also regulates mitochondrial biogenesis and transcription by directly phosphorylating peroxisome-proliferator-activated receptor γ coactivator 1α (PGC-1α), an important transcriptional coactivator (57). In worms, a direct ortholog of PGC-1α is not reported but MDT-15 acts as a functional ortholog (58). We assessed these molecule’s role using loss of function mutants in swimming exercise-mediated functional improvement (Fig. 3A). The absence of either *crh-1* or *hda-4* did not affect the enhancement in recovery index through swimming exercise (Fig S3A) ruling out the roles of CREB and HDAC5 in this process. However, the swimming-mediated improvement in behavioural recovery is not observed in *daf-16* and *mdt-15* mutants (Fig.3B, Fig.S3A). The basal level of axon regrowth in these mutants is equivalent to that of wild type (Fig. 3C-D, Fig.S3B). However, the enhanced regrowth length after swimming exercise is not found in both the *daf-16 (lf) and mdt-15(lf)* mutants unlike the wild type (Fig.3E). The swimming quality represented by thrashing frequency is comparable to the wild type for both groups (Fig.S3C). The absence of enhanced PTRI of aged *daf-16(lf)* animals following swimming, strengthened the role DAF-16 being a mediator of exercise’s beneficial effects. Impaired swimming is observed in the aged *mdt-15(lf)* animals resulting in exclusion from the paradigm. Additionally, The AICAR treatment also failed to enhance the recovery index in the *daf-16(lf)* and *mdt-15(lf)* indicating the roles of these transcriptional regulators downstream to AMPK (Fig. 3E). AMPK agonist treatment also does not enhance the PTRI in aged animals similar to axon regeneration (Fig. S3F). This result puts DAF-16 and MDT-15 as downstream effectors of AMPK in the context of swimming exercise.

**Figure 3.**
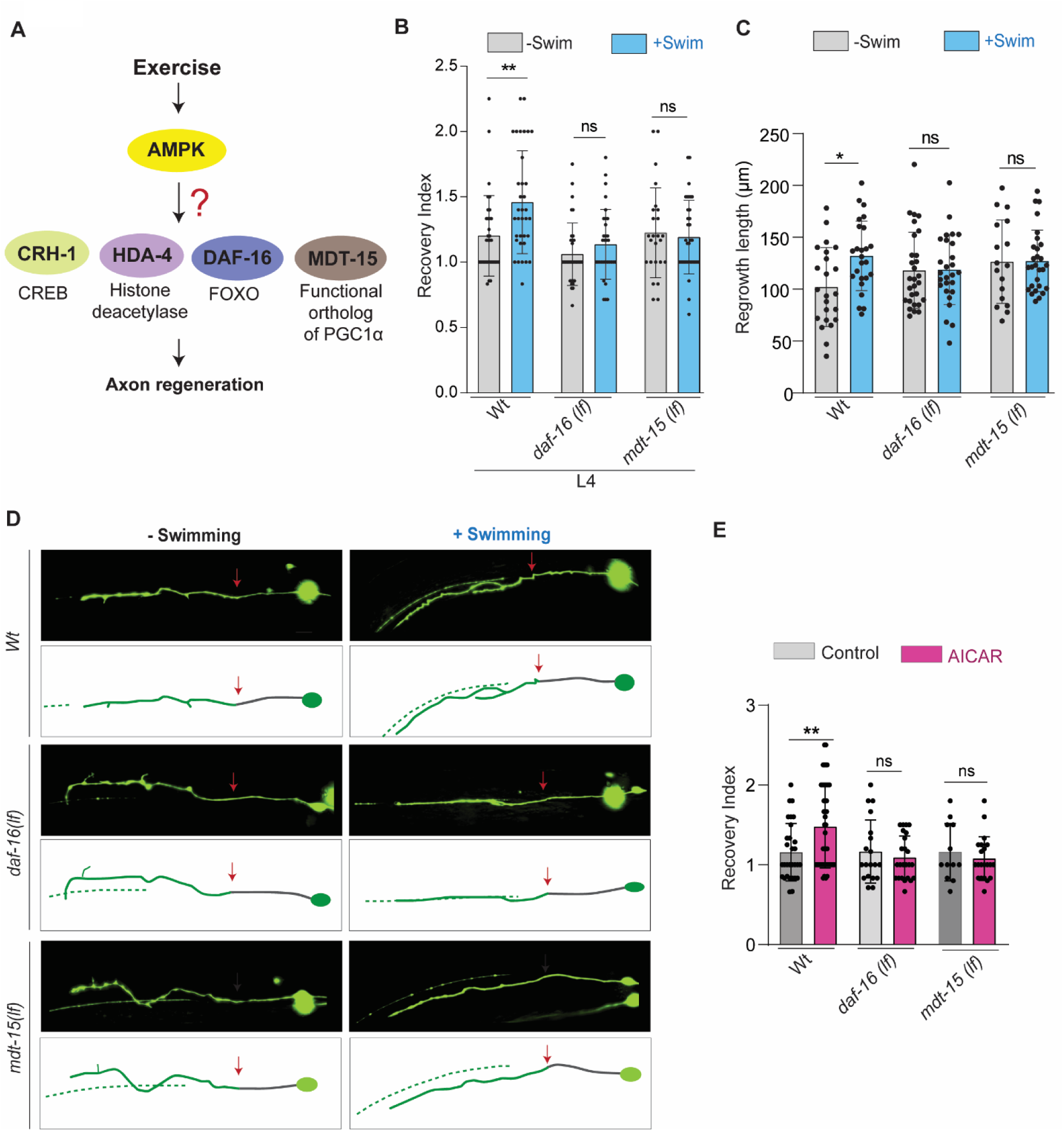
DAF-16 and MDT-15 are required for swimming-mediated enhanced functional recovery after PLM axon injury. **(A)** Schematics showing important downstream effectors of AMPK which may mediate the positive effects of swimming for functional improvement. **(B)** Bar graph showing functional recovery represented as recovery index for *daf-16 (lf)* and *mdt-15 (lf)* mutant worms after a 90-minute swimming session. N = 3-4 independent replicates, n = 24-44 number of worms tested. **(C)** Quantification of regrowth length at 24 h post axotomy as represented in D.N=3-4,n=17-29. **(D)** Microscopic image showing the axonal regrowth at 24 h post axotomy with or without swimming exercise for *daf-16(lf)* and *mdt-15 (lf)* worms. The red arrow shows the position of the axotomy. **(E)** AICAR (1mM) treatment does not improve the functional recovery of *daf-16(lf)* and *mdt-15(lf)* worms. N=3-4,n=12-45.Statistics, for B, C, E, *p < 0.05, **p < 0.01 ANOVA with Tukey’s multiple comparison test. Error bars represent SD; ns, not significant.

### DAF-16 is required in both muscles and neurons while MDT-15 is only in neurons for improved functional recovery after swimming exercise

Tissue-specific rescue experiments indicated that AMPK is required in both muscles and neurons for exercise-mediated improved functional recovery (10). To understand the tissue-specific roles of the transcription factors that act downstream of AMPK signalling in the swimming exercise, we used a muscle and neuron-specific RNAi of *daf-16* and *mdt-15* (59, 60). We did not observe an improvement in functional recovery after axon regeneration with swimming when muscle-specific RNAi was done for *aak-2* and *daf-16* unlike the vector control group (L4440) (Fig. 4B). Interestingly, knockdown of *mdt-15* in body wall muscle did not affect the enhanced functional recovery due to swimming (Fig.4B). Therefore, DAF-16 is required in body-wall muscle for swimming mediated enhanced functional recovery whereas MDT-15 is not. The ageing paradigm also complemented this finding (Fig.S4 A-B). Knockdown of the upstream kinase LKB1/PAR-4, the scaffold protein for PAR-4 axin/PRY-1 and AAK-2 in muscle did not improve the PTRI of old animals upon swimming session (Fig. S4B). The result showed that knocking down DAF-16 but not MDT-15 in muscle inhibits the swimming-mediated PTRI improvement (Fig.S4B).

**Figure 4.**
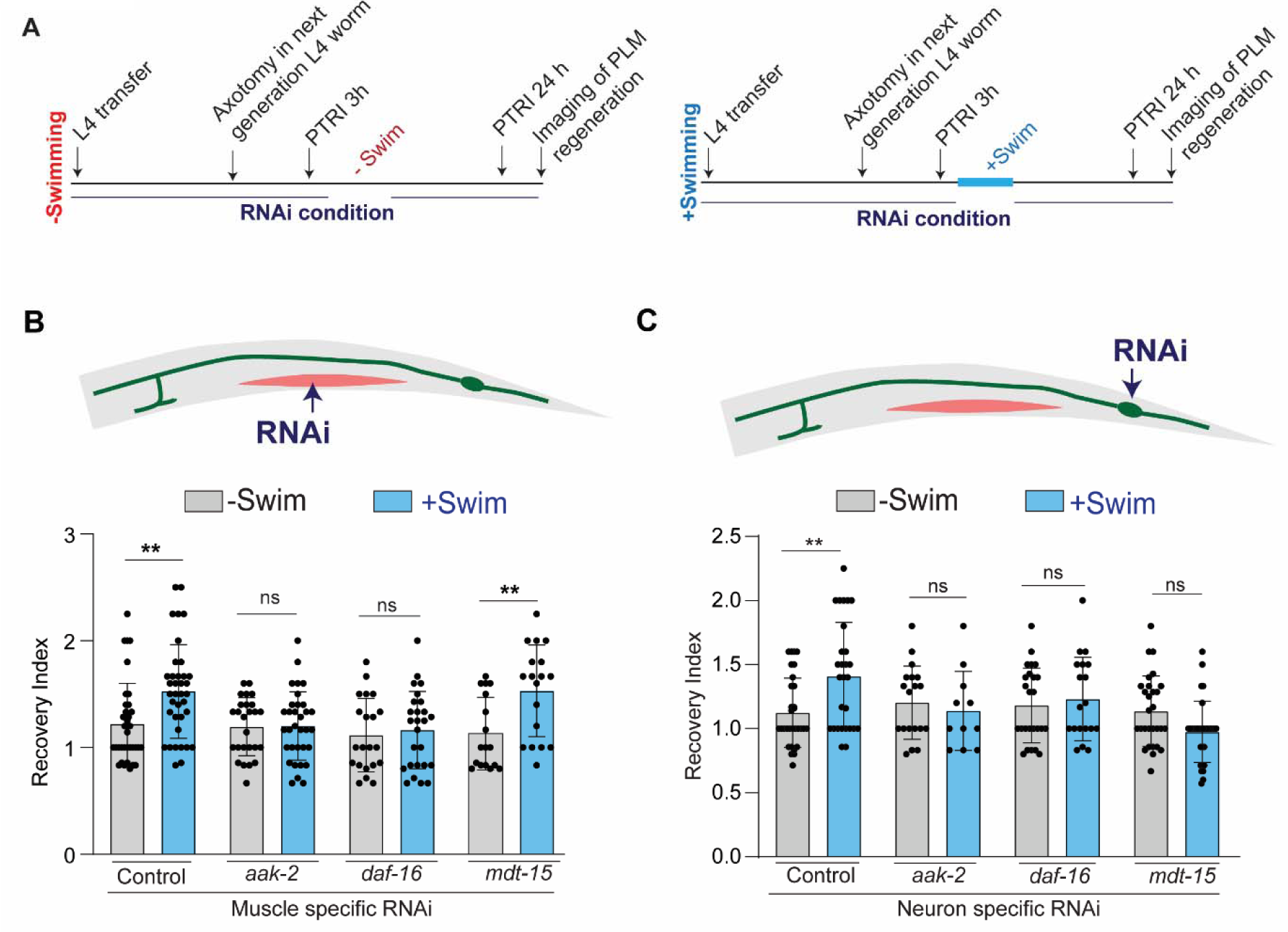
DAF-16 acts in body wall muscle and neuron whereas MDT-15 acts only in neuron for swimming-mediated enhancement in functional recovery after axon injury. **(A)** Paradigm schematics showing assessment of functional recovery after axotomy of L4 worms in RNAi condition. **(B)** Bar graph showing the recovery index for vector control (L4440), *aak-2*,*daf-16 and mdt-15* muscle-specific RNAi at 24 h postaxotomy. Axotomy was performed at the L4 stage. Muscle specific RNAi strain *rde-1(ne219); kzIs20* [(P*hlh-1*::*rde-1* + *sur-5*p::NLS::GFP); *tbIS222* (P*mec4*:: mcherry)]. N = 3 independent replicates, n = 17–37 number of worms tested **(C)** Bar graph showing the recovery index for vector control (L4440),*aak-2*,*daf-16* and *mdt-15* neuron-specific RNAi at 24 h postaxotomy. Axotomy was performed at the L4 stage. Neuron specific RNAi strain : [*sid-1(pk3321*) *him-5(e1490)*; l*in-15B(n744)*; *uIs72* (pCFJ90 (P*myo-2::mCherry*) + P*unc-119::sid-1* + P*mec-18::mec-18*::GFP)].N=3,n=11-31. Statistics, for A, D *p < 0.05, **p < 0.01 ANOVA with Tukey’s multiple comparison test. For B, C *p < 0.05 Unpaired t-test. Error bars represent SD; ns, not significant

Similarly, neuronal knockdown of both *aak-2* and *daf-16* hindered the improvement in recovery index through swimming session after axonal injury as compared to the control condition (Fig.4C). Similar observation was made upon neuronal RNAi of *mdt-15* (Fig.4C). It is clear from the bar graph that in neuronal knockdown of MDT-15 as well as DAF-16 impeded the enhanced functional recovery following swimming (Fig.4C). These RNAi experiments shows that DAF-16 has a differential roles across both neuronal and muscle tissues whereas MDT-15 has a neuron-specific role.

### Neuronal functions of both DAF-16 and MDT-15 are required for enhanced regrowth and ventral targeting due to swimming exercise

To understand what aspects of neuronal rewiring are controlled by these two downstream effectors, we further characterized the anatomical aspects of regrowth in muscle and neuron-specific RNAi conditions. We measured the regrowth length at early time points after axotomy in the tissue-specific knockdown. We found that the enhancement of axon regrowth at 10 h post-axotomy due to swimming exercise remained unaffected by the muscle-specific knockdown of either *daf-16* or *mdt-15* (Fig.5A). However, in contrast to muscle knockdown neuronal knockdown of each of *aak-2*, *daf-16* and *mdt-15* perturbs the enhanced regrowth mediated by the swimming (Fig.5B).

**Figure 5.**
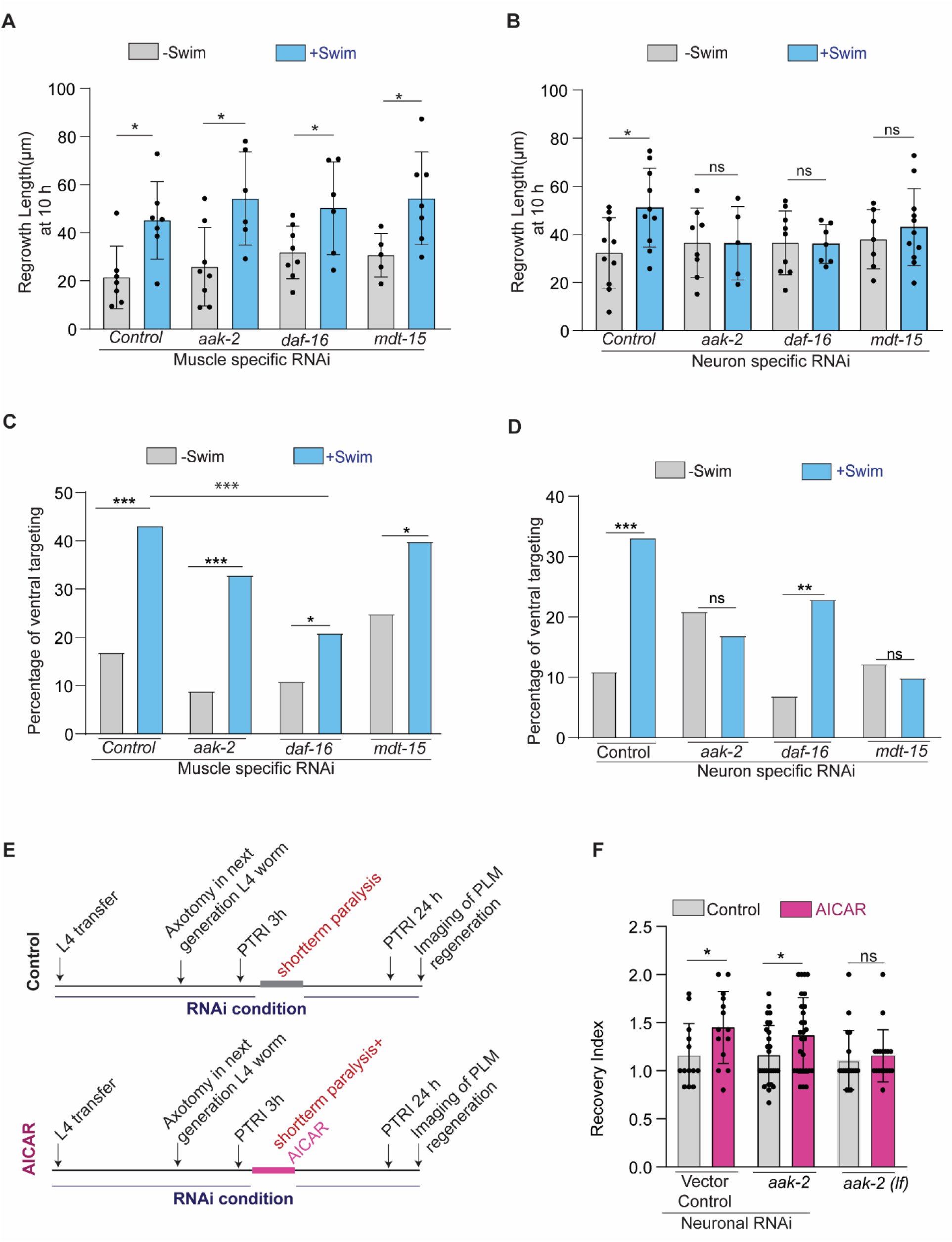
DAF-16 regulates axon regeneration from body wall muscle and neuron whereas MDT-15 from the neuron. **(A)** Regrowth length at 10h post axotomy in muscle-specific RNAi condition with or without a swimming session. N = 2-3 independent replicates, n = 5-8 number of worms tested **(B)** Regrowth length at 10h post axotomy in muscle-specific RNAi condition with or without a swimming session. N=2-3,n=5-11 **(C)** Percentage of Ventral targeting events in muscle-specific RNAi condition with or without a swimming session. Ventral targeting is the type of regeneration in which the regenerated PLM axon is targeted towards the ventral nerve cord where developmentally synaptic connection occurs. N=3,n>20 worms tested **(D)** Percentage of Ventral targeting events in neuron-specific RNAi condition with or without a swimming session. N=3,n>20 worms tested **(E)** The paradigm of AICAR treatment for the axotomized RNAi worms. **(F)** Recovery index of the AICAR treated neuronal aak-2 knockdown worms. N=3,n=13-28. Muscle specific RNAi strain *rde-1(ne219)*; *kzIs20* [(P*hlh-1::rde-1* + *sur-5p*::NLS::GFP); *tbIS222* (Pmec4:: mcherry)]. Neuron specific RNAi strain: [*sid-1(pk3321) him-5(e1490); lin-15B(n744); uIs72* (pCFJ90 (P*myo-2*::mCherry) + P*unc-119::sid-1* + P*mec-18::mec-18*::GFP)] Statistics for A, B, F *p < 0.05 Unpaired t-test. For C, D *p < 0.05 ***p < 0.001 Fisher’s exact test. Error bars represent SD; ns, not significant.

It was reported that when the regenerated axon is targeted and fasciculate along the ventral nerve cord it leads to functional recovery (40). These events were previously termed ventral targeting events and were enhanced as well by swimming exercise (10). The characterization of ventral targeting events in muscle-specific knockdown of *aak-2*,*daf-16* and *mdt-15* still showed a significant increase in ventral targeting similar to the control condition (Fig. 5C). However, in knockdown of *daf-16* affected the magnitude of enhancement in ventral targeting indicating that it regulates the ventral targeting enhancement from through a role in muscle.

In contrast, the neuronal knockdown of *aak-2* and *mdt-15* hampered the enhancement in ventral targeting following swimming (Fig.5D). In case of neuronal *daf-16*, still an enhancement in ventral targeting was observed (Fig. 5D). However, the magnitude of increase in ventral targeting is affected by neuron-specific knockdown of *aak-2*,*daf-16* and *mdt-15* (Fig.5D). These results showed that exercise mediated AMPK axis activation of *daf-16* regulates the ventral targeting from muscle and neuron whereas *mdt-15* regulates only from neuron to impart functional recovery.

We further characterized if non-neuronal activation of AMPK through drug treatment leads to enhanced behavioural recovery. For this we provided AICAR treatment to the neuronal *aak-2* knockdown worms after axotomy (Fig.5E). The AICAR treatment improved functional recovery in the neuronal *aak-2* knockdown condition (Fig.5F). This result acts as proof of principle that activation of non-neuronal AMPK can boost the functional recovery after axon regeneration.

### Swimming session promotes nuclear entry of DAF-16 through AMPK

To assay whether the swimming session directly affects DAF-16, we looked at the localization of the CRISPR-tagged endogenous DAF-16::mKate2 (61) before and after an exercise session. Usually DAF-16:: mKate2 is localized in many tissues including neurons in the nerve ring area (white arrows, Fig. 6A) as reported before. There is a nuclear pattern of localization throughout various tissues. We found that following a 90-minute swimming session, the nuclear localization pattern becomes prominent in various tissues including muscle cells as compared to the control condition (Yellow arrowheads, Fig 6A-B). The round-shaped nuclear pattern was prominent in the post-swimming group for the wild-type background, which was not seen in the *aak-2(lf)* background (Fig.6A-B). The nuclear localization is observed in several tissues including body wall muscle and intestine (Fig. 6B-C).

**Figure 6.**
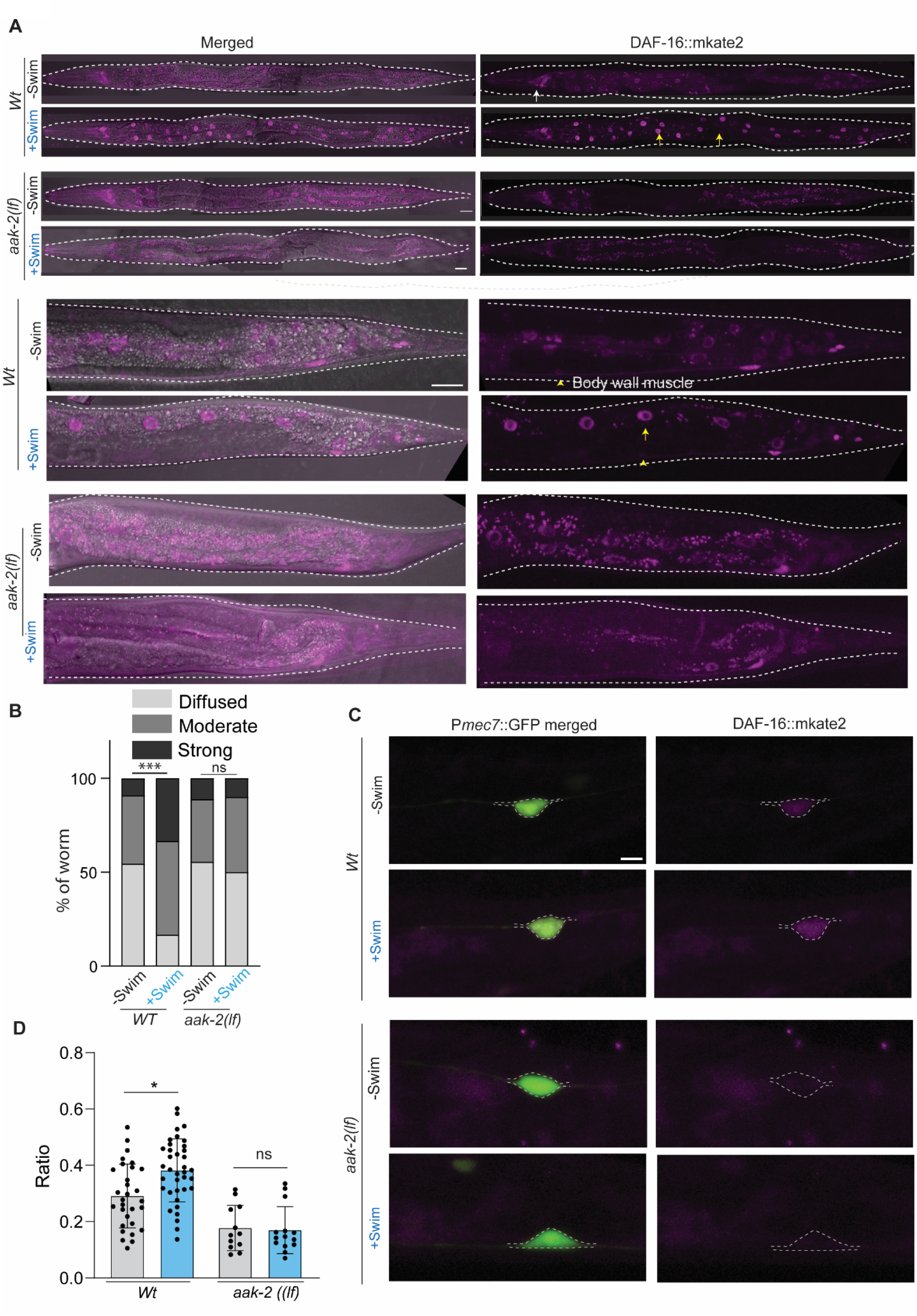
Swimming exercise induces DAF-16 nuclear localization and level mediated through AMPK. **(A-B)** Representative image showing enhanced DAF-16 localization after swimming exercise.DAF-16 localization is visualized by CRISPR tagged mkate2. The yellow arrow shows the intestine nucleus, yellow arrowhead shows the body wall muscle. The white arrow shows the nerve ring. **(C)** Quantification showing the percentage of worms and qualitative characterization of DAF-16 activation. Strong represents highly intense nuclear localization of DAF-16.Moderate means moderately intense DAF-16 nuclear localization and diffused means diffused pattern of DAF-16 level. N = 3 independent replicates, n >20 number of worms tested. **(D)**Representative images showing DAF-16 level after swimming exercise in the PLM neuron cell body. **(E)** Ratiometric quantification of mkate2 to GFP in the PLM neuron cell body, N=3, n=14-37. Statistics for B ***p<0.001 Fisher’s Exact test, for E *p < 0.05 Unpaired t-test. Error bars represent SD; ns, not significant.

We also assayed the effect of swimming exercise on the localization of DAF-16::mKate2 in the PLM neurone before and after the swimming session (Fig 6D). We noticed that following a swimming session, localization of DAF-16::mKate2 is increased in the cell body (dotted trace, Fig. 6D). Diffusible GFP expressed in touch neuron was used as a control to quantify the relative change in DAF-16::mKate2 localization in the cell body of PLM. We found that in the wild-type background, the ratio of DAF-16::mKate2 vs GFP is increased in the post-swimming group as compared to the control (Fig. 6E). Such increment was not observed in the aak-2 mutant (Fig.6D-E). This finding shows that DAF-16 is regulated by AMPK following swimming exercise through AMPK signalling.

## Discussion

In this study, we have shown that the metabolic energy sensor AMP Kinase is activated through Liver Kinase B-1/ PAR-4 during the swimming exercise, which in turn activates two signalling axes, one in neuron and the other in muscle tissues. The FOXO family transcription factor DAF-16 acts both in neuron and muscle, whereas PGCα/MDT-15 exclusively acts in neuron to mediate the beneficial effects of swimming exercise in functional rewiring of injured PLM axon (Fig.7). Excitingly, both of these signalling arms could be activated by AMPK agonist AICR as exercise mimetic. Our analysis indicates the neuronal roles of DAF-16 and MDT-15 in speeding up the regenerative response including enhanced regrowth from the injured axon stump and enhanced ventral targeting, which leads to behavioural recovery (40). The behavioural recovery after the axon regeneration response is a cumulative readout of the axon regrowth, circuit rewiring and neuromuscular response.

**Figure 7.**
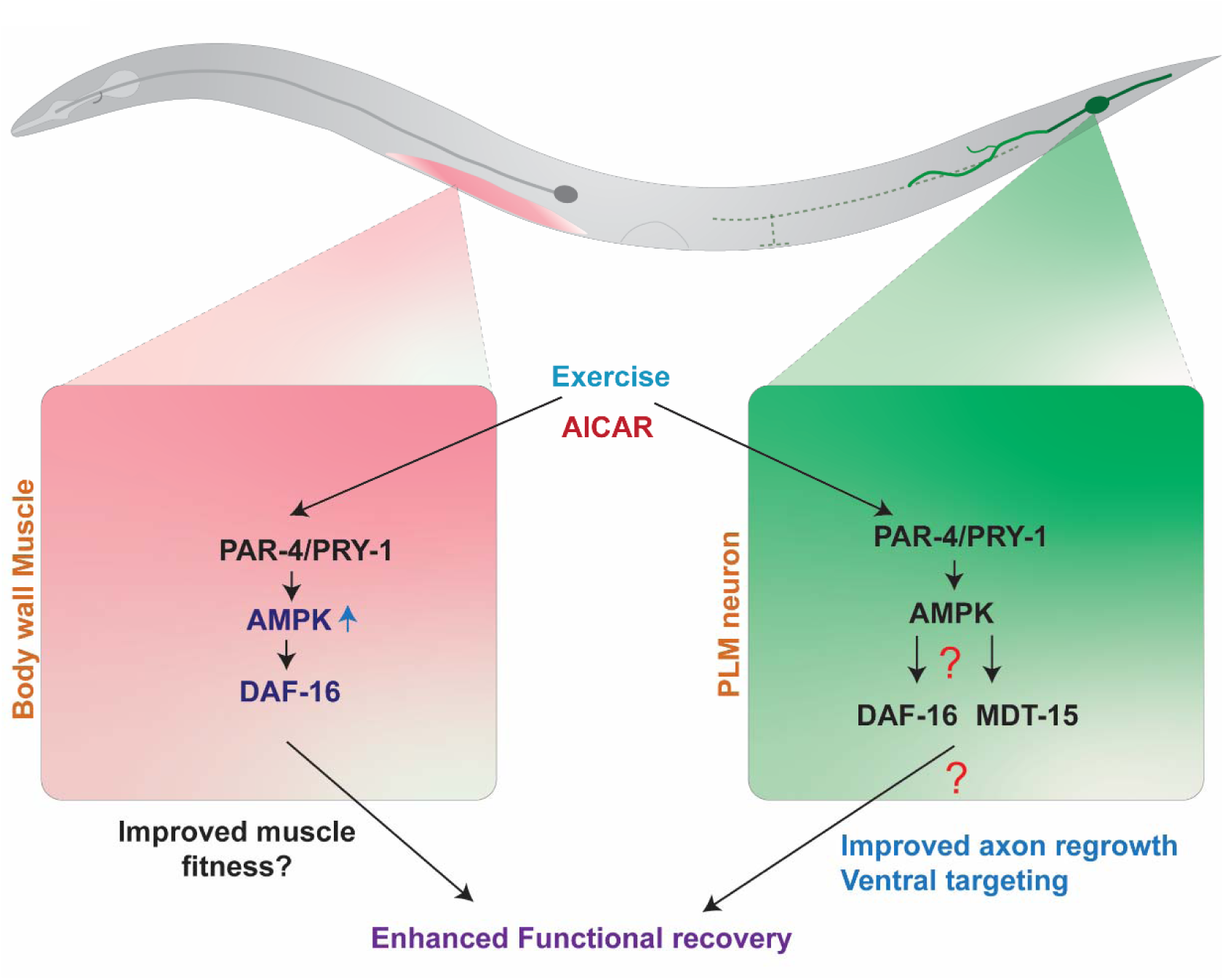
Schematics showing the summary of the AMPK signalling components required in body wall muscle and neuron for enhanced functional recovery after swimming exercise.

AMPK is a master regulator for maintaining cellular energetic balance by regulating ATP production and expenditure (23). AMPK activation after exercise has been linked to the regulation of system metabolism and health benefits (12). The finding that it is required for swimming exercise-mediated enhanced functional recovery after axon injury and averted functional decline due to ageing in *C.elegans* supports its conserved role (10). It is also required for swimming-mediated preservation of physical fitness during ageing (14). Immunoblot analysis of activated AMPK (p-AMPK) shows significantly high activation after swimming exercise in *C.elegans* (15). In this study, we found that acute activation of AMPK through AICAR treatment improved functional recovery after axon injury mimicking swimming exercise. In contrast, inhibition of AMPK activation through dorsomorphin blocked such enhancement by swimming session. This strengthens AMPK as a major regulatory hub for exercise-mediated beneficial effects. More interestingly, it puts AMPK activation through an AICAR-like agonist as a promising strategy to improve axon regeneration following an injury. AICAR treatment in mice leads to improvement in cognition and motor coordination both in the young and aged stages (62). AICAR treatment can be pursued further in other models for developing an axon regeneration improvement strategy.

AMPK activation is a highly regulated process that requires upstream factors including the kinases that phosphorylate the T172 position. Using a temperature-sensitive mutant of PAR-4/LKB-1, we found that it is required for swimming-associated improvement in both ageing and axon regeneration. Upregulation of LKB-1 in mouse CNS resulted in improved axon regeneration and enhanced locomotor functional recovery partly through AMPK (63). Our finding that PAR-4 is required for swimming-mediated recovery in posterior gentle touch response supports its axon regeneration-promoting role.

Activated AMPK regulates and restores energy balance by regulating metabolism acutely as well as initiating an adaptive response. AMPK regulates the activity of diverse downstream molecules regulated by direct phosphorylation (13). We tested selective downstream factors which are linked to exercise and axon regeneration. We found that DAF-16 (ortholog of FOXO) and MDT-15 (functional ortholog of PGC1α) are important downstream effectors for swimming-mediated enhancement in axon regeneration. Our result showed an increased nuclear localization of DAF-16 following swimming exercise both in neuronal and non-neuronal cells, which is dependent on the AMPK function. Previous reports also show increased DAF-16 localization in the nucleus after exercise (15, 64). Our finding along with this evidence supports a mechanistic model in which exercise-induced AMPK activation in both neuron and muscle leads to DAF-16 activation resulting in improved functional recovery. Swimming exercise improves the fitness of aged animals by active regulation of mitochondrial dynamics (Campos et al., 2023). In the absence of insulin receptor DAF-2 through DAF-16 transcription factor, aged animals preserve muscle mitochondrial morphology, and maintain higher muscle mitochondrial mass and high levels of intracellular ATP (65). Swimming exercise also induces autophagy in worms and DAF-16 is linked to this process (14, 15). Mitochondria and autophagy directly regulate axon regeneration potential after an injury in rodents and worm (66–68). AMPK regulates DAF-16 activity through direct phosphorylation at multiple sites (53).DLK-1 regulation by DAF-16 also affects axon regrowth following an injury (69). DAF-16 regulates many neuronal genes that affect cellular transport, axon guidance, and signalling, which can influence behavioural recovery (70).DAF-16 regulates netrin-mediated guidance of regenerating axons to the ventral nerve cord for successful behavioural recovery (40). Our data shows that DAF-16-mediated regulation of such targeting events contributes to enhanced recovery after exercise.

Our finding also shows that neuronal activity of MDT-15 leads to improved functional recovery after neuronal injury after swimming exercise. MDT-15 acts as a functional orthologue of PGC1α in *C.elegans* (58). MDT-15 encodes a subunit of the transcriptional coregulatory Mediator complex and regulates fatty acid metabolism genes, fasting-induced gene expression, xenobiotic and heavy metal detoxification genes and oxidative stress response (58, 71–76). Understanding how MDT-15 regulates functional recovery after axon regeneration is an interesting direction to explore further.

We also observed that when AMPK is knocked down in neuron, AICAR treatment could boost the non-neuronal function of AMPK to promote axon regeneration. This opens up an exciting window of opportunity to integrate the multiple signalling arms of AMPK signalling across various tissues.

## Materials & Methods

### Genetics and Caenorhabditis elegans strains

Standard conditions were followed to grow and maintain the strains using Nematode Growth Media (NGM) at 20° C (77). For some sets of experiments, worms were maintained at alternative temperatures and mentioned in the respective places. Standard *E. coli* OP50 strain was used for growing the worms. Detailed genotypes of the *C.elegans* strains used are denoted in Table S1. Standard nomenclature notion was followed to describe genotypes of the *C.elegans* strains used in this study. For example, *dlk-1(lf)* denotes that a loss of function allele i.e. *tm4024* was used in which DLK-1 is not functional. These genetic strains were acquired from the Caenorhabditis Genetics Centre (CGC) and genotyped using an appropriate strategy involving PCR, RFLP and sequencing.

### Synchronization of worms

Fifty to seventy gravid worms were transferred to freshly NGM plates containing OP50 bacteria and the worms were allowed to lay eggs for 2 hrs. After 2 hrs, the adults were removed from the plate and embryos were grown till the L4 stage. Then 60-70 L4 worms were transferred to fresh OP50 seeded NGM plates containing 50 μm 5-Fluoro deoxyuridine (FUDR; Sigma; catalog #F0503) (37). The worms were transferred to a new FuDR plate after 3 days. The adult day was counted from the time of L4 stage worm transfer for example after 24 h from the time of transfer, the worms are labelled as day 1 adults (A1 stage). For the experiments in this study, several life-stage worms were used and specified in subsequent places.

### Swimming exercise paradigm

We used a single session of swimming exercise in our study which is considered as an acute physical exercise paradigm (28, 29). For a swimming session, the worms were subjected to 1x M9 buffer in a 96-flat bottom plate for 90 minutes. A single worm was kept in a single well containing 200 µl buffer for the swimming session (swimming well). The plate was gently shaken at a 10-minute interval to avoid the settlement of worms. After the swimming session was over, the worms were recovered on NGM plates using a Pasteur pipette. The control no swimming group worms were kept in an unseeded NGM plate for the duration of the swimming session and were transferred to OP50 containing NGM plates after 90 minutes. We primarily used a 90-minute swimming session for our experiments as the ATP levels significantly dropped after this and worms switched to an episodic mode of swimming as per previous reports (10, 64, 78).

### Laser-assisted axotomy

Appropriate stage worms were immobilized on a 5% agarose bed prepared on top of a glass slide. Worms were physically immobilized in a 0.1-μm polystyrene bead solution (Polysciences® 00876-15) while sandwiched between the agarose and a coverslip. We axotomized only a single PLM neuron axon for all the experiments at a 50-60 µm distance from the cell body as described previously (37). The gap created by two laser pulses was around 7-10 µm. The “cut side” and the “control side” are the sides that correspond to the side of axotomized PLM neuron and uninjured PLM neuron respectively. We used a Bruker® system with a Spectra Physics® femtosecond multiphoton laser set-up for performing simultaneous imaging and axotomy as described previously (37). The system uses two Spectra Physics (Mai Tai with Deepsee) automated depression-compensated femtosecond lasers which are tunable in the wavelength range of 690-1020 nm. The GFP-labelled axons were imaged with a 920 nm laser while axotomy was performed using a 720 nm laser micropulse. For mcherry labelled PLM neuron axotomy, imaging was performed with a 1020 nm laser. A 3-mm galvanometer system was used for axotomy, while a 6-mm galvanometer scanning system was used for the imaging.

### Gentle touch assay

Gentle touch assay was performed for functional assessment after axon injury and in the context of ageing according to the previously published method (37, 79, 80). We gave ten alternative anterior and posterior gentle touch with the fine tip of an eye-lash and recorded the reversal response. We scored the response by giving the anterior touch to the forward-moving worm and the posterior touch to a backward-moving worm. The anterior touch was given in the area between the pharynx and vulva whereas the posterior touch was given between vulva and anus. The positive reversal response was denoted as 1 and 0 for the corresponding no response. The posterior touch response index (PTRI) was calculated by taking the ratio of total positive response out of the total touch given which is 10 in our case as reported before (37).

### Correlation of functional recovery with axon regeneration at a single neuronal level

After 3h of axotomy, each worm was kept on a single NGM plate and PTRI was assessed from each side of the worm. Then, worms were kept at 20 °C for 24 h followed by PTRI measurement. After the 24 h PTRI was assessed, the worms were transferred to an agarose bed glass slide and the individual worm was immobilized in a drop of 10mM levamisole solution. The anatomical regeneration pattern was assessed in an epi-fluorescence microscope (Nikkon Ti2 eclipse) or confocal microscope (Nikkon-A1 confocal system). The functional recovery was correlated with axon regeneration using the ‘Recovery Index’. The recovery index was calculated by the ratio of PTRI at 24 h to PTRI at 3h for a single side representing the touch response of a single PLM neuron.

### Imaging of regenerated PLM axons

The regenerated PLM axons were imaged after 24 hours of axotomy. The worms were immobilized on a 5% agarose pad using a 10 mM levamisole solution. The imaging was done using a Nikkon Ti2 epi-fluorescence microscope or a Nikkon A1 confocal system.

In the epi-fluorescence system, PLM axons were imaged under a 40x oil (NA-1.1) objective using a Zyla EMCCD camera at the exposure value of 200 ms. GFP-labelled PLM axons were imaged using the appropriate filter set and z-scanning with a step size of 0.5 µm.

In the confocal system, PLM axons were imaged using a 40X air (NA-0.9) objective. The GFP labelled axons were imaged using a 488 nm solid-state laser powered on 20%. The PMT power for the 488-nm channel was 100 with an offset value set at 20.PLM axons were imaged at a z-scanning step size of 0.5 µm. For the mcherry labelled axons, imaging was done using a 561 nm laser power to 20%. The PMT power for the 561 nm channel was 110 with an offset value at 15. A similar 0.5 µm z-scanning step was used. The regrowth length was calculated from the z-stacked image using Image J with the plugin simple neurite tracer.

### AICAR treatment on paralyzed worms

The animals were given a short exposure (∼15 seconds) to 5 mM of levamisole hydrochloride to temporarily paralyse them after 3h post axotomy PTRI was assessed. This brief levamisole treatment was effective in perturbing swimming during the 90-minute swimming session in the swimming well. This short levamisole treatment does not affect gentle touch response (10). Therefore, we used this paradigm to test the effect of AICAR treatment on functional axon regeneration. These paralyzed worms were subjected to a swimming well containing the specified concentration of AICAR (Sigma-Aldrich; catalogue #A9978) in 1x M9 buffer. For the control groups, paralyzed worms were kept only in 1x M9 buffer for the duration of the swimming session. After the session, worms were recovered to NGM plates containing OP50 bacteria. For the ageing paradigm, AICAR treatment was done similarly to the specified aged animals.

### Thrashing frequency analysis of swimming worms

We recorded videos of swimming worms using a Leica MC 120 HD camera for the thrashing frequency analysis. We recorded for 30 s at 15 frames/s using LAS V4.4 software at 1.25× magnification in the camera attached to a Leica stereo microscope M165 FC. We calculated body bends per second from the video analysis by using the ImageJ software plugin wrMTrck (http://www.phage.dk/plugins/wrmtrck.html) and converted the values to thrashing frequency (Thrashes/minute). One body bend was defined as a bending direction change at the mid-body section (10)

### Tissue specific RNAi condition

We used a similar induction method for efficient RNAi as reported earlier (81). We re-streaked the required gene from the library stored at -80° C on an LB agar plate containing 50 µg/ml carbenicillin and 12.5 µg/ml tetracycline and followed by overnight incubation at 37°C. The bacteria was restreaked again before the inoculation into liquid culture. The bacteria were grown on 4 ml LB broth containing 50 µg/ml carbenicillin and 12.5 µg/ml tetracycline till the OD_600_ reached 0.7-0.8 value. Then the culture was centrifuged at 4300 g for 2 minutes to harvest the bacteria and the supernatant was thoroughly dried. The pellet was resuspended in 1 ml of sterilized 1x M9 containing 1.5 mM IPTG, 50 µg/ml carbenicillin and 12.5 µg/ml tetracycline. This resuspended culture was seeded to the previously prepared NGM plates containing the same concentration of IPTG, carbenicillin and tetracycline. Approximately 330 µl of resuspended culture was seeded onto a 60 mm NGM plate whereas 50 µl was seeded onto a 35 mm NGM plate. For preparing these NGM plates, after pouring the NGM with the abovementioned chemicals the plates were allowed 24 h room temperature solidification followed by 24 h incubation at 4 °C. The seeded bacteria culture was dried in the laminar hood for 2 h and then incubated at 25 °C for 36 hours. After the incubation was over, 7-10 L4 worms of the required genotype were transferred to a 60 mm plate and kept at 20°C for growth. In each batch control genes (*dhc-1 and ama-1*) were used to ensure the induction and optimum RNAi condition. We used the L4 progenies from the RNAi plates for the experiments. For muscle-specific RNAi, we used NR350 strain which is developed based on muscle-specific rescue of RDE-1 using *hlh-1* promoter in *rde-1(lf)* background and is an established strain for this purpose (59). For neuron-specific RNAi, we used the TU3595 strain which is reported to be optimized for efficient RNAi in gentle touch neurons (60).

### Ageing of RNAi worms

We picked L4 worms from the RNAi plates described in the above section and transferred them to RNAi bacteria-seeded NGM plates containing the previously mentioned concentration of carbenicillin, tetracycline, IPTG and also contained FUDR (50 mM). The worms were aged to the required adulthood stage in such plates and worms were transferred to new plates at 3 days intervals. At the specific adulthood stage, worms were picked and subjected to swimming and non-swimming condition. Following swimming worms were recovered in RNAi plates and after 24h functional assessment was performed.

### Axotomy and functional recovery assessment of RNAi worms

The PLM neuron axotomy was performed with worms picked from the RNAi plates according to the above-mentioned procedure. After axotomy, worms were recovered on RNAi bacterial plates and kept for 3 hours for recovery from the stress of the axotomy procedure. Then we separated each axotomized worm into a 35mm RNAi bacterial plate and assessed the gentle touch behaviour as per the method described previously. After the behavioural assessment, a swimming session was introduced to a group of worms while the control group did not undergo a swimming session. Following the swimming session, worms were recovered back onto the RNAi plates and kept in the incubator for 24 hours for axon regeneration. After 24 h, again gentle touch response assay was performed to assess the functional recovery and then worms were subjected to the microscopic analysis for anatomical characterization of axon regeneration.

### Imaging of DAF-16 localization pattern after swimming

We used the mkate2 tagged DAF-16 reporter strain to analyse the effect of swimming exercise on DAF-16 localization (61). A single worm was mounted on a 5% agarose pad using 10mM levamisole and imaged within 10 minutes using a Nikon AXR confocal microscope. The conditions were maintained to minimize the external stress. The control worms were kept in the unseeded plate for 90 minutes while the swimming group underwent a swimming session in the well.

### Statistics

We used GraphPad Prism software version 9.0.2 for all the statistical analysis in this study. We used an unpaired t test with Welch’s correction for two-way comparison. We used Fisher’s exact test for comparing proportions and ANOVA (nonparametric) with a post hoc Tukey’s multiple comparisons test for comparing three or more groups. Details of the tests are described in the figure legends. The values represent mean ± standard deviation. Each bar of the graph represents sample numbers (n) accumulated over the total number of biological replicates (N) in a given experiment.

## Acknowledgements

We thank NBRP, Japan and Caenorhabditis Genetics Center (CGC) for strains. CGC is supported by the NIH Office of Research Infrastructure Programs (P40 OD010440). We thank Shirshendu Dey and Swagata Dey for maintaining the 2-photon and the micro point UV laser set-up respectively. We thank Kavinila Selvarasu for the help in RNAi experiments. This work is supported by the NBRC core fund from the Department of Biotechnology, The India Alliance DBT Wellcome Senior fellowship (Grant #IA/S/22/1/506243) to Anindya Ghosh-Roy and a grant from Science and Engineering Research Board (SERB: CRG/2019/002194) to Anindya Ghosh-Roy.

## Author Contributions

Sibaram Behera and Anindya Ghosh-Roy designed experiments. Sibaram Behera generated mutant combinations using genetic crosses, performed all the experiments experiments and analyzed data. Sibaram Behera, and Anindya Ghosh-Roy wrote the manuscript.

## References

1. Maas AIR, et al. (2017) Traumatic brain injury: integrated approaches to improve prevention, clinical care, and research. Lancet Neurol 16(12):987–1048.

2. Injury GBDTB & Spinal Cord Injury C (2019) Global, regional, and national burden of traumatic brain injury and spinal cord injury, 1990-2016: a systematic analysis for the Global Burden of Disease Study 2016. Lancet Neurol 18(1):56–87.

3. Zheng B & Tuszynski MH (2023) Regulation of axonal regeneration after mammalian spinal cord injury. Nat Rev Mol Cell Biol 24(6):396–413.

4. He Z & Jin Y (2016) Intrinsic Control of Axon Regeneration. Neuron 90(3):437–451.

5. Griffin JM, et al. (2023) Rehabilitation enhances epothilone-induced locomotor recovery after spinal cord injury. Brain Commun 5(1):fcad005.

6. van Praag H (2008) Neurogenesis and exercise: past and future directions. Neuromolecular Med 10(2):128–140.

7. Ruegsegger GN & Booth FW (2018) Health Benefits of Exercise. Cold Spring Harb Perspect Med 8(7).

8. Pedersen BK (2019) Physical activity and muscle-brain crosstalk. Nat Rev Endocrinol 15(7):383–392.

9. Molteni R, Zheng JQ, Ying Z, Gomez-Pinilla F, & Twiss JL (2004) Voluntary exercise increases axonal regeneration from sensory neurons. Proc Natl Acad Sci U S A 101(22):8473–8478.

10. Kumar S, Behera S, Basu A, Dey S, & Ghosh-Roy A (2021) Swimming Exercise Promotes Post-injury Axon Regeneration and Functional Restoration through AMPK. eNeuro 8(3).

11. Maugeri G, et al. (2021) The role of exercise on peripheral nerve regeneration: from animal model to clinical application. Heliyon 7(11):e08281.

12. Spaulding HR & Yan Z (2022) AMPK and the Adaptation to Exercise. Annu Rev Physiol 84:209–227.

13. Steinberg GR & Hardie DG (2023) New insights into activation and function of the AMPK. Nat Rev Mol Cell Biol 24(4):255–272.

14. Campos JC, et al. (2023) Exercise preserves physical fitness during aging through AMPK and mitochondrial dynamics. Proc Natl Acad Sci U S A 120(2):e2204750120.

15. Chen YL, et al. (2023) Physical exercise attenuates age-related muscle atrophy and exhibits anti-ageing effects via the adiponectin receptor 1 signalling. J Cachexia Sarcopenia Muscle.

16. Hardie DG, Schaffer BE, & Brunet A (2016) AMPK: An Energy-Sensing Pathway with Multiple Inputs and Outputs. Trends Cell Biol 26(3):190–201.

17. Hawley SA, et al. (2003) Complexes between the LKB1 tumor suppressor, STRAD alpha/beta and MO25 alpha/beta are upstream kinases in the AMP-activated protein kinase cascade. J Biol 2(4):28.

18. Woods A, et al. (2003) LKB1 is the upstream kinase in the AMP-activated protein kinase cascade. Curr Biol 13(22):2004–2008.

19. Shaw RJ, et al. (2004) The tumor suppressor LKB1 kinase directly activates AMP-activated kinase and regulates apoptosis in response to energy stress. Proc Natl Acad Sci U S A 101(10):3329–3335.

20. Hawley SA, et al. (2005) Calmodulin-dependent protein kinase kinase-beta is an alternative upstream kinase for AMP-activated protein kinase. Cell Metab 2(1):9–19.

21. Woods A, et al. (2005) Ca2+/calmodulin-dependent protein kinase kinase-beta acts upstream of AMP-activated protein kinase in mammalian cells. Cell Metab 2(1):21–33.

22. Mihaylova MM & Shaw RJ (2011) The AMPK signalling pathway coordinates cell growth, autophagy and metabolism. Nat Cell Biol 13(9):1016–1023.

23. Herzig S & Shaw RJ (2018) AMPK: guardian of metabolism and mitochondrial homeostasis. Nat Rev Mol Cell Biol 19(2):121–135.

24. Steinberg GR & Carling D (2019) AMP-activated protein kinase: the current landscape for drug development. Nat Rev Drug Discov 18(7):527–551.

25. Jeong JH, et al. (2023) A new AMPK isoform mediates glucose-restriction induced longevity non-cell autonomously by promoting membrane fluidity. Nat Commun 14(1):288.

26. Yanik MF, et al. (2004) Neurosurgery: functional regeneration after laser axotomy. Nature 432(7019):822.

27. Byrne AB & Hammarlund M (2017) Axon regeneration in C. elegans: Worming our way to mechanisms of axon regeneration. Exp Neurol 287(Pt 3):300–309.

28. Laranjeiro R, Harinath G, Burke D, Braeckman BP, & Driscoll M (2017) Single swim sessions in C. elegans induce key features of mammalian exercise. BMC Biol 15(1):30.

29. Laranjeiro R, et al. (2019) Swim exercise in Caenorhabditis elegans extends neuromuscular and gut healthspan, enhances learning ability, and protects against neurodegeneration. Proc Natl Acad Sci U S A 116(47):23829–23839.

30. Hisamoto N & Matsumoto K (2017) Signal transduction cascades in axon regeneration: insights from C. elegans. Curr Opin Genet Dev 44:54–60.

31. Ghosh-Roy A, Wu Z, Goncharov A, Jin Y, & Chisholm AD (2010) Calcium and cyclic AMP promote axonal regeneration in Caenorhabditis elegans and require DLK-1 kinase. J Neurosci 30(9):3175–3183.

32. Hammarlund M, Nix P, Hauth L, Jorgensen EM, & Bastiani M (2009) Axon regeneration requires a conserved MAP kinase pathway. Science 323(5915):802–806.

33. Yan D, Wu Z, Chisholm AD, & Jin Y (2009) The DLK-1 kinase promotes mRNA stability and local translation in C. elegans synapses and axon regeneration. Cell 138(5):1005–1018.

34. Zou Y, et al. (2013) Developmental decline in neuronal regeneration by the progressive change of two intrinsic timers. Science 340(6130):372–376.

35. Wu Z, et al. (2007) Caenorhabditis elegans neuronal regeneration is influenced by life stage, ephrin signaling, and synaptic branching. Proc Natl Acad Sci U S A 104(38):15132–15137.

36. El Bejjani R & Hammarlund M (2012) Notch signaling inhibits axon regeneration. Neuron 73(2):268–278.

37. Basu A, et al. (2017) let-7 miRNA controls CED-7 homotypic adhesion and EFF-1-mediated axonal self-fusion to restore touch sensation following injury. Proc Natl Acad Sci U S A 114(47):E10206–E10215.

38. Abay ZC, et al. (2017) Phosphatidylserine save-me signals drive functional recovery of severed axons in Caenorhabditis elegans. Proc Natl Acad Sci U S A 114(47):E10196–E10205.

39. Ding C & Hammarlund M (2018) Aberrant information transfer interferes with functional axon regeneration. Elife 7.

40. Basu A, Behera S, Bhardwaj S, Dey S, & Ghosh-Roy A (2021) Regulation of UNC-40/DCC and UNC-6/Netrin by DAF-16 promotes functional rewiring of the injured axon. Development 148(11).

41. Corton JM, Gillespie JG, Hawley SA, & Hardie DG (1995) 5-aminoimidazole-4-carboxamide ribonucleoside. A specific method for activating AMP-activated protein kinase in intact cells? Eur J Biochem 229(2):558–565.

42. Kim J, Yang G, Kim Y, Kim J, & Ha J (2016) AMPK activators: mechanisms of action and physiological activities. Exp Mol Med 48(4):e224.

43. Ju S, et al. (2022) C. elegans monitor energy status via the AMPK pathway to trigger innate immune responses against bacterial pathogens. Commun Biol 5(1):643.

44. Burkewitz K, et al. (2015) Neuronal CRTC-1 governs systemic mitochondrial metabolism and lifespan via a catecholamine signal. Cell 160(5):842–855.

45. Zhou G, et al. (2001) Role of AMP-activated protein kinase in mechanism of metformin action. J Clin Invest 108(8):1167–1174.

46. Berry BJ, Baldzizhar A, Nieves TO, & Wojtovich AP (2020) Neuronal AMPK coordinates mitochondrial energy sensing and hypoxia resistance in C. elegans. FASEB J 34(12):16333–16347.

47. Zhang YL, et al. (2013) AMP as a low-energy charge signal autonomously initiates assembly of AXIN-AMPK-LKB1 complex for AMPK activation. Cell Metab 18(4):546–555.

48. Chen J, et al. (2017) Metformin extends C. elegans lifespan through lysosomal pathway. Elife 6.

49. Mallick A, Ranawade A, van den Berg W, & Gupta BP (2020) Axin-Mediated Regulation of Lifespan and Muscle Health in C. elegans Requires AMPK-FOXO Signaling. iScience 23(12):101843.

50. Trefts E & Shaw RJ (2021) AMPK: restoring metabolic homeostasis over space and time. Mol Cell 81(18):3677–3690.

51. Canto C & Auwerx J (2010) AMP-activated protein kinase and its downstream transcriptional pathways. Cell Mol Life Sci 67(20):3407–3423.

52. Greer EL, et al. (2007) The energy sensor AMP-activated protein kinase directly regulates the mammalian FOXO3 transcription factor. J Biol Chem 282(41):30107–30119.

53. Greer EL, et al. (2007) An AMPK-FOXO pathway mediates longevity induced by a novel method of dietary restriction in C. elegans. Curr Biol 17(19):1646–1656.

54. Thomson DM, et al. (2008) AMP-activated protein kinase phosphorylates transcription factors of the CREB family. J Appl Physiol (1985) 104(2):429–438.

55. McGee SL, et al. (2008) AMP-activated protein kinase regulates GLUT4 transcription by phosphorylating histone deacetylase 5. Diabetes 57(4):860–867.

56. Cho Y, Sloutsky R, Naegle KM, & Cavalli V (2013) Injury-induced HDAC5 nuclear export is essential for axon regeneration. Cell 155(4):894–908.

57. Jager S, Handschin C, St-Pierre J, & Spiegelman BM (2007) AMP-activated protein kinase (AMPK) action in skeletal muscle via direct phosphorylation of PGC-1alpha. Proc Natl Acad Sci U S A 104(29):12017–12022.

58. Taubert S, Van Gilst MR, Hansen M, & Yamamoto KR (2006) A Mediator subunit, MDT-15, integrates regulation of fatty acid metabolism by NHR-49-dependent and -independent pathways in C. elegans. Genes Dev 20(9):1137–1149.

59. Qadota H, et al. (2007) Establishment of a tissue-specific RNAi system in C. elegans. Gene 400(1-2):166–173.

60. Calixto A, Chelur D, Topalidou I, Chen X, & Chalfie M (2010) Enhanced neuronal RNAi in C. elegans using SID-1. Nat Methods 7(7):554–559.

61. Aghayeva U, Bhattacharya A, & Hobert O (2020) A panel of fluorophore-tagged daf-16 alleles. MicroPubl Biol 2020.

62. Kobilo T, et al. (2014) AMPK agonist AICAR improves cognition and motor coordination in young and aged mice. Learn Mem 21(2):119–126.

63. Ohtake Y, et al. (2019) Promoting Axon Regeneration in Adult CNS by Targeting Liver Kinase B1. Mol Ther 27(1):102–117.

64. Chaudhari SN & Kipreos ET (2017) Increased mitochondrial fusion allows the survival of older animals in diverse C. elegans longevity pathways. Nat Commun 8(1):182.

65. Wang H, Webster P, Chen L, & Fisher AL (2019) Cell-autonomous and non-autonomous roles of daf-16 in muscle function and mitochondrial capacity in aging C. elegans. Aging (Albany NY) 11(8):2295–2311.

66. Han SM, Baig HS, & Hammarlund M (2016) Mitochondria Localize to Injured Axons to Support Regeneration. Neuron 92(6):1308–1323.

67. Cartoni R, et al. (2016) The Mammalian-Specific Protein Armcx1 Regulates Mitochondrial Transport during Axon Regeneration. Neuron 92(6):1294–1307.

68. Ko SH, Apple EC, Liu Z, & Chen L (2020) Age-dependent autophagy induction after injury promotes axon regeneration by limiting NOTCH. Autophagy 16(11):2052–2068.

69. Byrne AB, et al. (2014) Insulin/IGF1 signaling inhibits age-dependent axon regeneration. Neuron 81(3):561–573.

70. Kaletsky R, et al. (2016) The C. elegans adult neuronal IIS/FOXO transcriptome reveals adult phenotype regulators. Nature 529(7584):92–96.

71. Lee D, et al. (2019) MDT-15/MED15 permits longevity at low temperature via enhancing lipidostasis and proteostasis. PLoS Biol 17(8):e3000415.

72. Yang F, et al. (2006) An ARC/Mediator subunit required for SREBP control of cholesterol and lipid homeostasis. Nature 442(7103):700–704.

73. Taubert S, Hansen M, Van Gilst MR, Cooper SB, & Yamamoto KR (2008) The Mediator subunit MDT-15 confers metabolic adaptation to ingested material. PLoS Genet 4(2):e1000021.

74. Shomer N, et al. (2019) Mediator subunit MDT-15/MED15 and Nuclear Receptor HIZR-1/HNF4 cooperate to regulate toxic metal stress responses in Caenorhabditis elegans. PLoS Genet 15(12):e1008508.

75. Mao K, et al. (2019) Mitochondrial Dysfunction in C. elegans Activates Mitochondrial Relocalization and Nuclear Hormone Receptor-Dependent Detoxification Genes. Cell Metab 29(5):1182–1191 e1184.

76. Goh GY, et al. (2014) The conserved Mediator subunit MDT-15 is required for oxidative stress responses in Caenorhabditis elegans. Aging Cell 13(1):70–79.

77. Brenner S (1974) The genetics of Caenorhabditis elegans. Genetics 77(1):71–94.

78. Ghosh R & Emmons SW (2008) Episodic swimming behavior in the nematode C. elegans. J Exp Biol 211(Pt 23):3703–3711.

79. Chalfie M & Sulston J (1981) Developmental genetics of the mechanosensory neurons of Caenorhabditis elegans. Dev Biol 82(2):358–370.

80. Chalfie M, et al. (1985) The neural circuit for touch sensitivity in Caenorhabditis elegans. J Neurosci 5(4):956–964.

81. Kamath RS, Martinez-Campos M, Zipperlen P, Fraser AG, & Ahringer J (2001) Effectiveness of specific RNA-mediated interference through ingested double-stranded RNA in Caenorhabditis elegans. Genome Biol 2(1):RESEARCH0002.

