## Supporting File for "Exercise-induced differential transcriptional output of AMPK signalling improves axon regeneration and functional recovery"

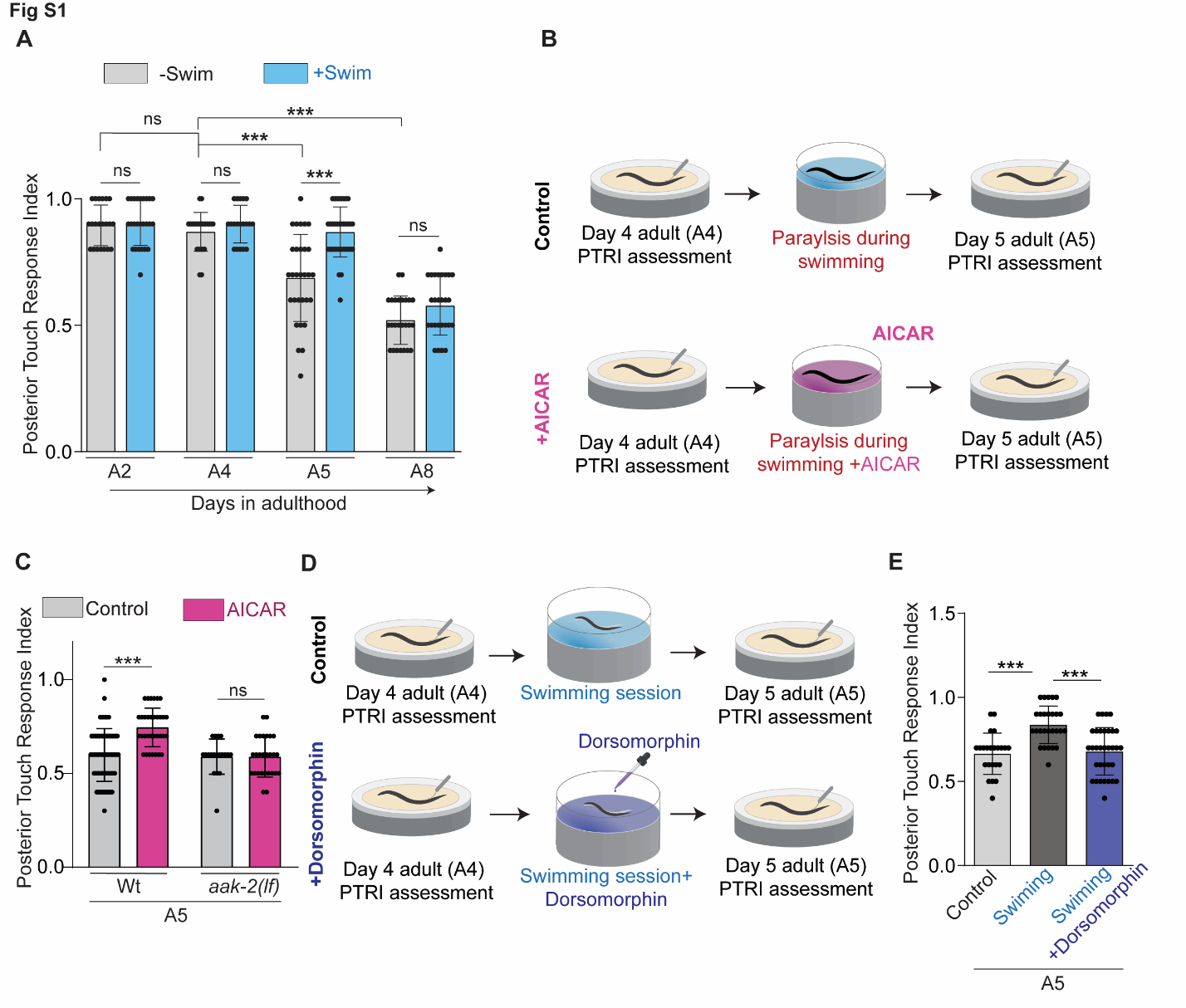

Figure S1 Acute activation of AMPK through AICAR treatment ameliorates age dependent decline in posterior touch response. (A) Bar graph showing Posterior Touch Response Index (PTRI) for A5 worms with or without swimming session at A4 adult stage .N = 3-5 independent replicates, n = 20-52 number of worms. (B) Schematics showing paradigm to test the effect of AICAR treatment on the ageing associated posterior touch response decline. (C) Bar graph showing Posterior Touch Response Index (PTRI) for A5 worms with or without AICAR treatment at A4 adult stage. N = 3-5 ,n = 20-52. (D) Schematics of the paradigm to test the effect of dorsomorphin treatment on the ageing associated posterior touch response decline. (E) Bar graph showing the effect of dorsomorphin treatment during swimming exercise on PTRI of A5 worms. N=3-4,n=18-24. Statistics, for A,C,E ***p < 0.001 ANOVA with Tukey’s multiple comparison test. Error bars represent SD; ns, not significant.

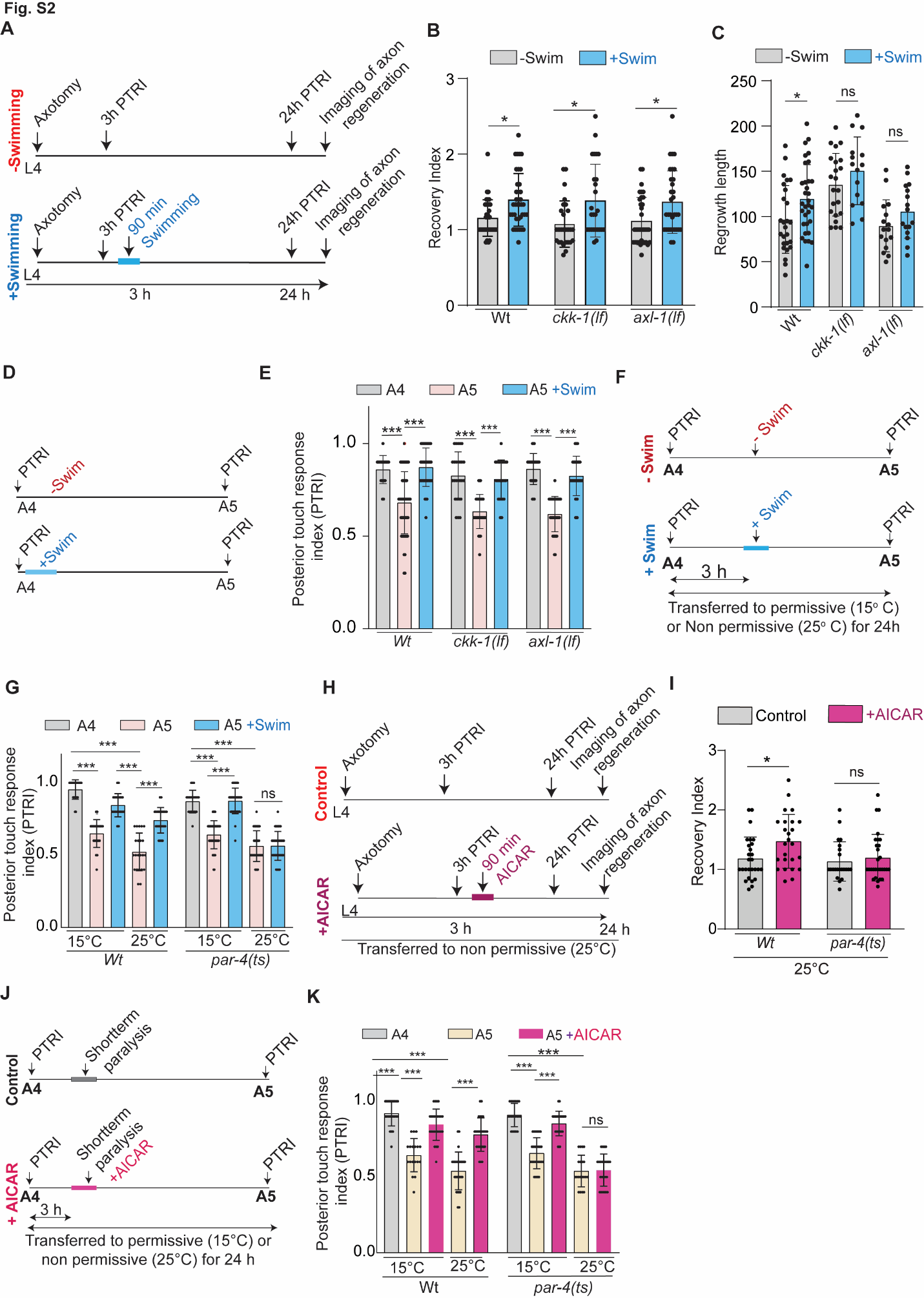
 Figure S2 Swimming-mediated prevention of ageing-associated decline in posterior gentle touch response requires AMPK upstream kinase liver kinase B1/PAR-4. (A) Schematics showing paradigm to test role upstream mediators of AMPK utilizing the readout of functional recovery in axon regeneration after a single 90-minute swimming session. (B) Bar graph showing recovery index at L4 stage axotomy of *ckk-1(lf)* and *axl-1(lf)* worms after swimming exercise. N = 3-5 independent replicates, n = 20-52 number of worms. (C) Regrowth length at 24 h post axotomy for the mentioned genotypes.(D) Paradigm to test effect of swimming on aging associated decline for *ckk-1(lf)* and *axl-1 (lf)* worms.(E) PTRI values at day 5 stage with swimming as depicted in panel D for the mentioned genotypes. (F) Paradigm using temperature-sensitive allele of PAR-4 to test its involvement in swimming-mediated prevention of decline in PTRI with ageing. The worms were aged till the A4 stage at a permissive temperature (15 ºC) and then divided into two groups. The permissive group was maintained at 15ºC while the non-permissive group was maintained at 25 ºC afterwards. The worms were maintained for 3 hours at the respective temperatures before a 90-minute swimming session was introduced. (G) Bar graph showing the PTRI values obtained for wild type and *par-4(ts)* worms at permissive (15 ºC) and non-permissive (25 ºC) temperatures after swimming exercise. N=3-4, n=21-45.(H) Schematics showing AICAR treatment paradigm on *par-4(ts)* mutant worms.(I) Recovery index of *par-4(ts)* animals following AICAR treatment. N=3,n=21-29. (J) Paradigm to test the effect of 1mM AICAR treatment to directly activate AMPK of temporarily paralyzed wildtype and *par-4(ts)* worms on PTRI improvement at A5 stage. (K) Bar graph showing the PTRI values obtained for wild type and *par-4(ts)* worms at permissive (15 ºC) and non-permissive (25 ºC) temperature after 1mM AICAR treatment. N=2-4, n=18-27.tatistics, for B,E, G,I and K ***p < 0.001 ANOVA with Tukey’s multiple comparison test. For C, Unpaired t-test *p < 0.05 Error bars represent SD; ns, not significant.

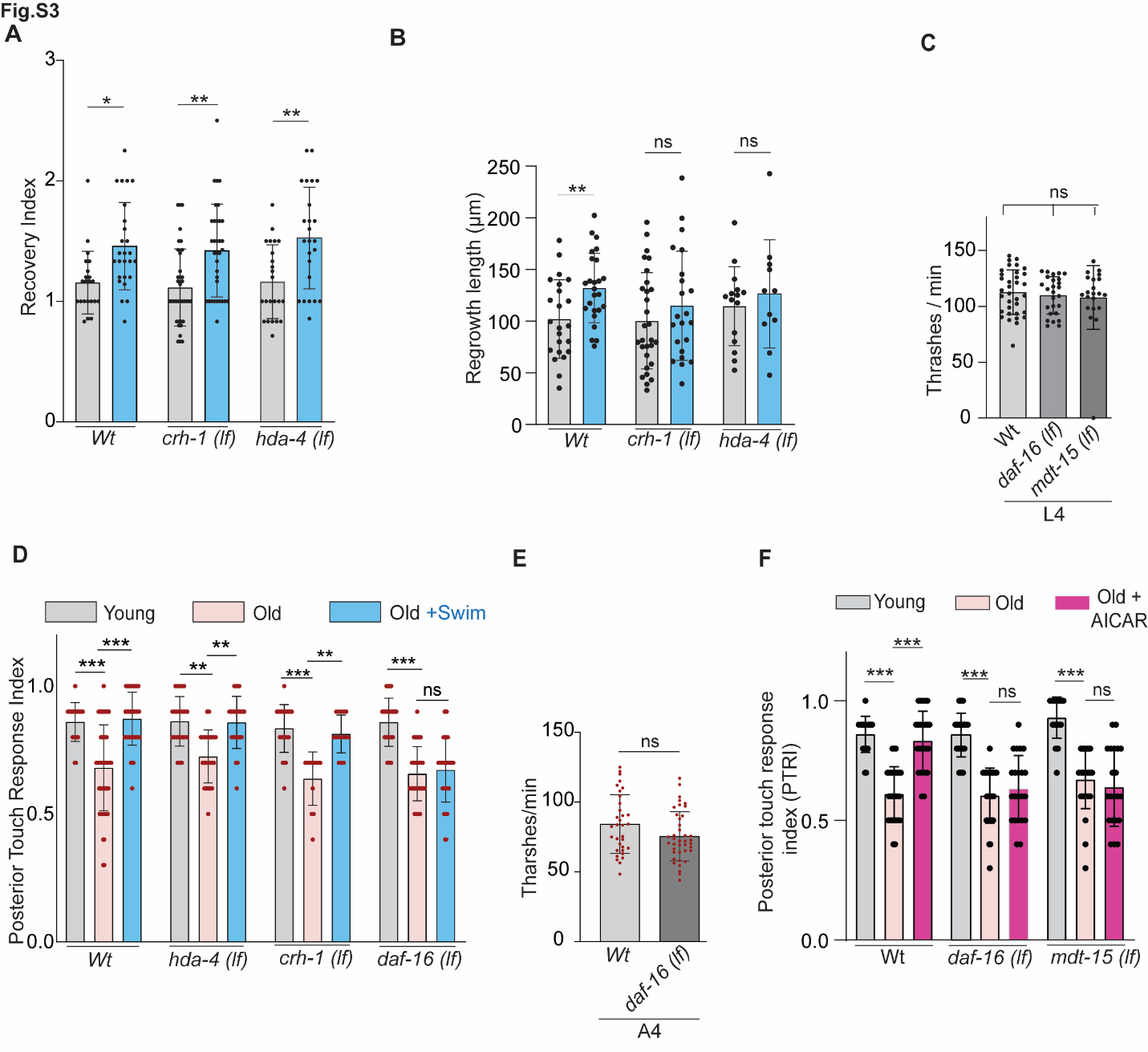

Figure S3 Role of DAF-16 and MDT-15 in swimming exercise mediated improvement of posterior gentle touch response in aged worms. (A) Recovery index values after swimming exercise for *crh-1(lf)* and *had-4(lf)* animals. Axotomy was performed at L4 stage. N = 2-4 independent replicates, n = 15–52 number of worms tested (B) Regrowth length at 24 h postaxotomy with swimming exercise for mentioned genotypes at L4 stage axon injury. N=3, n=20-28. (C) Bar graph showing thrashing frequency comparison of wildtype, daf-16(lf) and mdt-15(lf) worms at L4 stage. N=3,n=35-42. (D) Observed PTRI of aged worms after the introduction of a 90-minute swimming exercise 24 hours before the PTRI assessment. For *wildtype* and *daf-16(lf),* young represents the A4 stage and old represents the A5 stage*.* For *had-4(lf)* and *crh-1(lf)*, young indicates the A2 stage and Old indicates the A3 stage. N = 2-4 independent replicates, n = 15–52 number of worms tested. (E) Bar graph showing the thrashing frequency of A4 stage wildtype and *daf-16(lf)* worms. N=3,n=33-40 (F) Bar graph showing the observed PTRI values after 1 mM AICAR treatment to the aged *daf-16(lf)* and *mdt-15(lf)* worms. For wildtype and *daf-16(lf)*, young represents the A4 stage and old represents the A5 stage. For *mdt-15(lf)*, young indicates the A2 stage and Old indicates the A3 stage. N=3,n=20-40. Statistics, for B, E, **p < 0.01, ***p < 0.001 ANOVA with Tukey’s multiple comparison test. For C, unpaired t-test Error bars represent SD; ns, not significant.

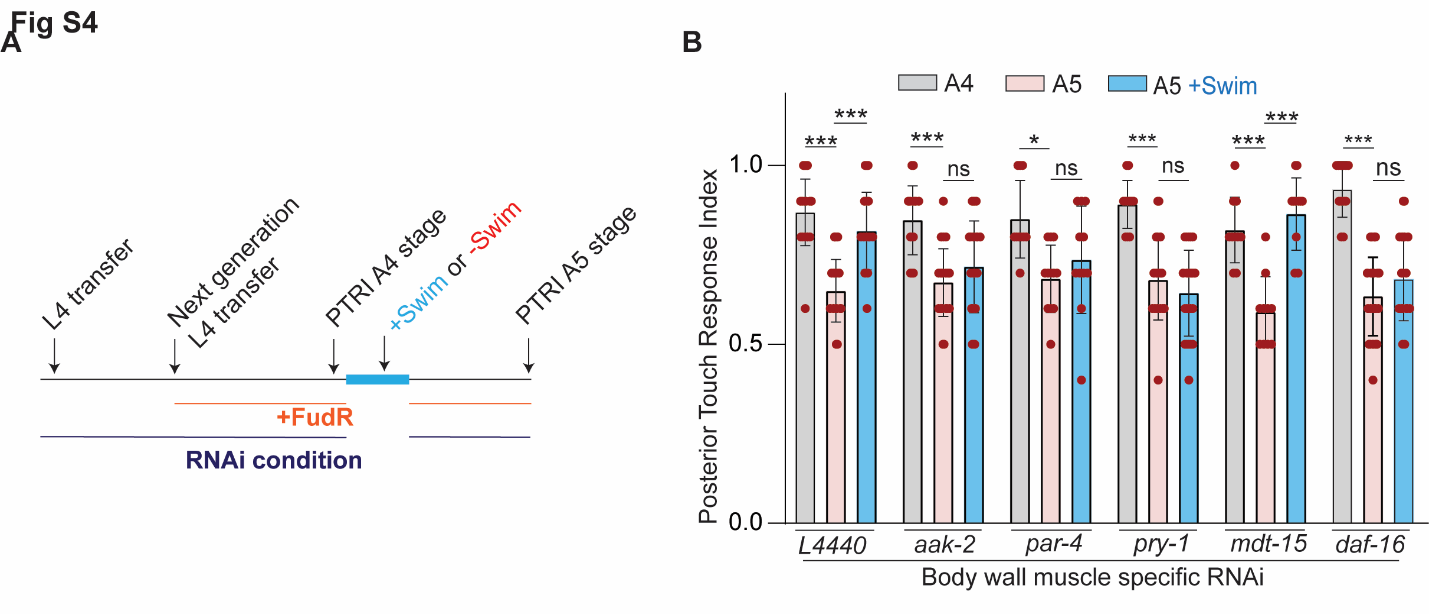

**Figure S4** Muscle-specific knockdown of AMPK signalling components inhibit the improved posterior touch response after swimming exercise**. (A)** Schematics showing the aging paradigm in RNAi condition to test the effect of swimming on PTRI drop. **(F)** Bar plot showing the PTRI values after swimming in muscle-specific knockdown of AMPK signalling components. Muscle specific RNAi strain *rde-1(ne219*) ; *kzIs20* [(P*hlh-1*::*rde-1* + *sur-5*p::NLS::GFP).N=3-5, n=10-36. Statistics, for B, *p < 0.05, **p < 0.01, ***p < 0.001 unpaired t-test Error bars represent SD; ns, not significant.

**Table S1** Details of the strains used in the study

| **Strain Number** | **Genotype** | **Source** |
| --- | --- | --- |
| N2 | *Wild type* | CGC |
| NBR270 | *muIs32 [Pmec-7:GFP+ lin-15(+)] II* | AGR lab |
| NBR1036 | *aak-1(tm1944) III; muIs32 II* | AGRL |
| RB754 | *aak-2(ok524) X* | CGC |
| NBR590 | *aak-2(ok524)X; muIs32 II* | AGRL |
| VC691 | *ckk-1(ok1033) III* | CGC |
| NBR1037 | *ckk-1(ok1033) III; muIs32 II* | AGRL |
| KN611 | *axl-1(tm1095) I* | CGC |
| NBR376 | *axl-1(tm1095) I; muIs32 II* | AGRL |
| KK300 | *par-4(it57) V* | CGC |
| NBR327 | *par-4(it57) V; muIs32 II* | AGRL |
| RB758 | *hda-4(ok518) X* | CGC |
| NBR 16 | *hda-4(ok518) X; muIs32 II* | AGRL |
| YT17 | *crh-1(tz2) III* | CGC |
| NBR1029 | *crh-1(tz2) III,muIS32 II* | AGRL |
| XA7702 | *mdt-15 (tm2182) III* | CGC |
| NBR105 | *mdt-15 (tm2182) III; muIs32 II* | AGRL |
| CF1038 | *daf-16 (mu86) I* | CGC |
| NBR541 | *daf-16 (mu86) I; muIs32 II* | AGRL |
| NR350 | *rde-1 (ne219); kzIs20 (hlh-1p::rde-1 + sur-5p::NLS::GFP)* | CGC |
| NBR 521 | *tbIs222 (Pmec-4:: mcherry);rde-1(ne219);kzIs20* | AGRL |
| TU3595 | *sid-1(pk3321) him-5(e1490) V; lin-15B(n744) X; uIs72 [pCFJ90(myo-2p::mCherry) + unc-119p::sid-1 + mec-18p::mec-18::GFP]* | CGC |
| OH13908 | *daf-16(ot821[daf-16::mKate2::3xFLAG]) I* | CGC |
| NBR1087 | *Ot821;muIS32* | AGRL |
